# Lactation Remodels the Maternal Lung to Promote Preemptive Innate Defense

**DOI:** 10.64898/2026.10.02.756233

**Authors:** Thomas R Cafiero, Soo Ching Lee, Chenyan Huang, Krist H Antunes Fernandes, Raju Shivarathri, Dede Setiawan, Harsha Sen, Jessica Ribeiro de Souza, Leafy Ye, Ricardo Mallarino, Fenna M Krienen, Jigar V. Desai, Ai Ing Lim

**Author notes:** Corresponding author: Ai Ing Lim.

## Abstract

Lactation is a hallmark of mammalian evolution, imposing extraordinary physiological demands on mothers to nourish their offspring. Yet how mothers maintain their own health while meeting these demands remains poorly understood. Here, we discover that lactation establishes a preemptive innate defense program in the maternal lung through enhanced epithelial production of surfactant proteins and mucin, vascular remodeling that supports granulocyte recruitment and accumulation of activated intravascular neutrophils. These tissue-wide adaptations safeguard lactating dams against diverse respiratory pathogens. We further identify prolactin and CGRP as neuroendocrine signals contributing to pulmonary remodeling and uncover parallel lactation-associated granulocyte adaptations in marsupial and non-human primate models. Together, our findings expand the framework of reproductive immunology, revealing that reproduction couples systemic fetal tolerance with tissue-specific immune adaptations that safeguard maternal health during lactation.

## Introduction

Lactation is one of the most energetically demanding physiological states in mammalian life (*1*). Mothers undergo profound adaptations across multiple organ systems to produce nutrient- and immune factor-rich breastmilk. The remarkable ability to nourish offspring without compromising maternal health is fundamental to the survival of both the mother and child, ultimately shaping species fitness across generations. Yet, despite lactation being a defining hallmark of mammals, surprisingly little is known about how it reshapes maternal immunity beyond the mammary gland (*2*–*7*). Research and clinical care often shift toward the newborn after delivery, leaving the question of how maternal immunity adapts during lactation largely unexplored.

The prevailing view that reproduction compromises maternal immune defense to promote fetal tolerance largely stems from studies of systemic and placental immunity during pregnancy, which documented a shift toward type 2 responses and expansion of regulatory T cells (*8*–*12*). In parallel with such systemic immune modulations, we recently discovered that reproduction progressively remodels the maternal intestinal immune-epithelial axis, culminating during lactation in enhanced innate defense against diverse enteric pathogens (*13*). Whether such adaptations represent an intestine-specific phenomenon or a broader physiological strategy that enhances maternal immune defense across barrier tissues remains unknown.

The lung is a vital barrier tissue that performs the essential function of gas exchange while continuously exposed to numerous airborne pathogens. Effective pulmonary defense relies on the coordinated actions of epithelial cells that form physical and antimicrobial barriers, endothelial cells that regulate leukocyte recruitment, and immune cells that eliminate invading pathogens (*14*–*18*). Here, we show that lactation promotes coordinated epithelial, endothelial and immune remodeling in the maternal lung, establishing a tissue-wide preemptive innate defense program that protects mothers against diverse respiratory pathogens. We further identify prolactin and CGRP as key neuroendocrine signals contributing to this lactation-specific immune adaptation.

## Results

### Maternal lung epithelium and endothelium harbor heightened innate defense programs during lactation

To investigate whether lactation remodels the maternal lung, we first examined the epithelial and endothelial compartments, which are central to barrier defense and immune cell recruitment. We performed unbiased transcriptomic profiling of purified cells from age-matched nulliparous controls and dams at day 12 of lactation, corresponding to the period of maximal breastmilk production (**Fig. 1A, fig. S1A**) (*19*). In the absence of infection, the lung epithelium underwent extensive transcriptional alteration during lactation, with 1922 genes significantly upregulated (**Fig. 1B, Table S1**). Many of the upregulated genes encoded components of epithelial innate defense programs, including surfactants (*Sftpa1, Sftpb, Sftpc, Sftpd*), complement components (*C1ql2, C4bp, C5ar2, C6, Hc*), mucins (*Muc1, Muc4, Mucl3, Muc20*), lysozymes (*Lyz1, Lyz2*), and mediators of immune signaling (*Csf2, Cxcl15, Cxcl17*) (**Fig. 1C**). Strikingly, the transcriptomic induction was also observed at the protein level. The lungs of lactating dams exhibited significantly elevated surfactant proteins A and D, collectins that bind and neutralize inhaled pathogens (*20*–*22*), together with MUC1, a membrane-bound mucin that reinforces epithelial barrier function and mucociliary clearance (*23*–*25*). In addition, the protein level of granulocyte colony-stimulating factor (G-CSF), a cytokine that promotes granulocyte production and survival, was significantly increased in the lungs of lactating dams (**Fig. 1D**).

**Figure 1.**
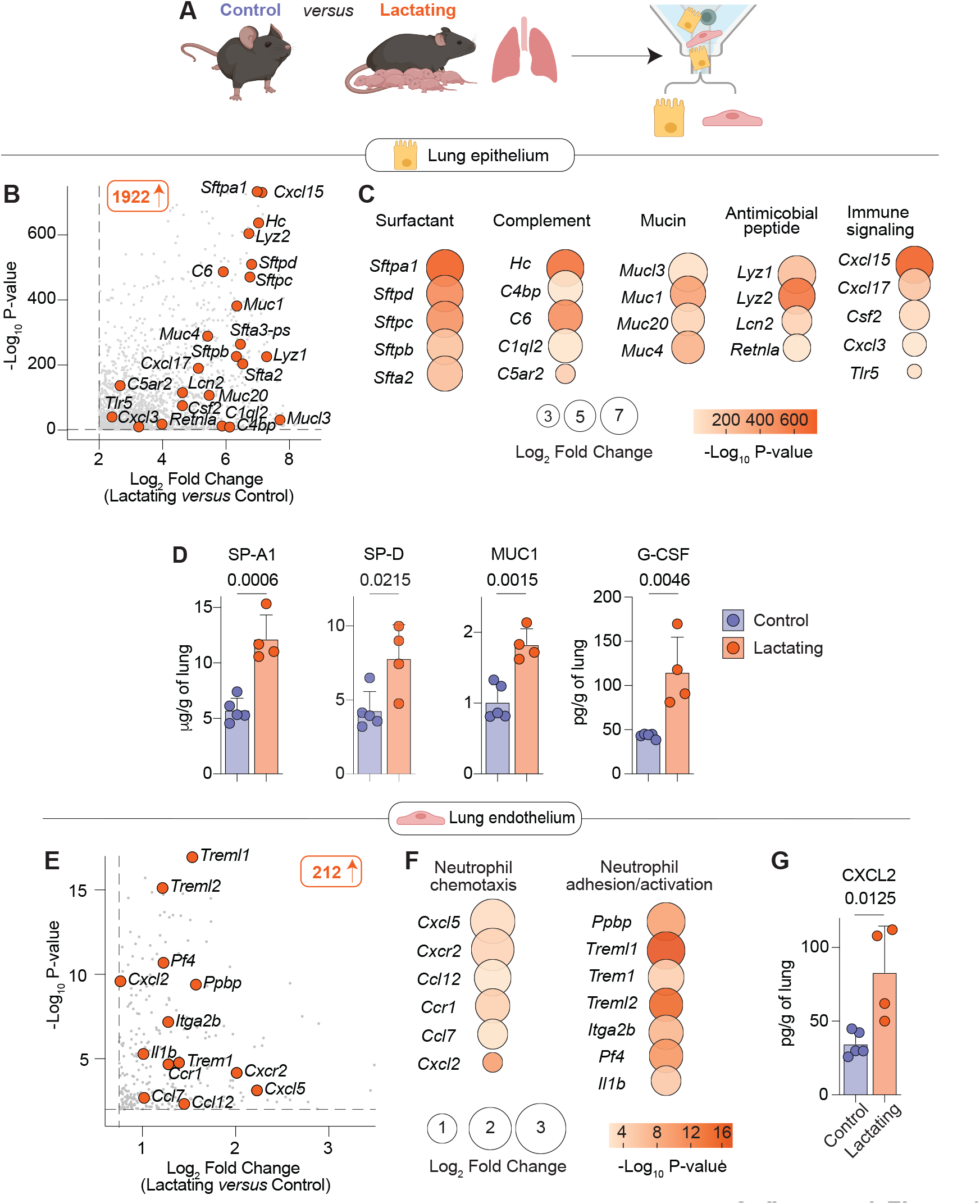
Maternal lung epithelial and endothelial cells enhance innate defense programs during lactation. **(A)**Lung epithelium and endothelium were isolated by FACS from day 12 lactating dams and age-matched nulliparous controls for transcriptomic profiling. (**B**) Volcano plot of differentially overexpressed genes in lung epithelial cells from lactating versus nulliparous mice. (**C**) Fold changes and *P* values of epithelial innate defense genes in lactating versus nulliparous mice. (**D**) Protein concentrations in lung measured by ELISA. (**E**) Volcano plot of differentially overexpressed genes in lung endothelial cells from lactating versus nulliparous mice. (**F**) Fold changes and *P* values of neutrophil recruitment and activation genes in lung endothelial cells from lactating mice versus nulliparous controls. (**G**) Protein concentration in lung measured by ELISA. Data in (D) and (G) represent mean ± SD and were analyzed by unpaired t-test. Each dot represents one mouse. Data are representative of two independent experiments. *P* values are indicated.

In parallel with epithelial activation, the lung endothelium also underwent transcriptional changes during lactation, although less extensive than that observed in the epithelium, with 212 genes significantly upregulated, including genes involved in neutrophil chemotaxis (*Cxcl2, Cxcl5, Cxcr2, Ccl12*), activation or adhesion (*Trem1, Treml2, Treml1, Itga2b*) (**Fig. 1E, F, Table S2**). Consistent with these transcriptomic changes, the potent neutrophil chemoattractant, CXCL2, was also elevated at the protein level in the lungs of lactating dams (**Fig. 1G**).

Together, these findings demonstrate that during lactation, even in the absence of pathogen exposure, the maternal lung undergoes extensive remodeling toward a preemptive innate defense program, characterized by enhanced epithelial barrier functions with an endothelium poised for granulocyte recruitment.

### Activated neutrophils accumulate in the lungs during lactation

The coordinated induction of signals promoting granulopoiesis and granulocyte recruitment during lactation led us to ask whether these changes were accompanied by alterations in pulmonary immune cell composition. Multiparameter flow cytometric analysis revealed that the pulmonary immune compartment remained largely unchanged during lactation, with comparable numbers of total immune cells and most innate and adaptive populations relative to age-matched nulliparous controls (**fig. S2A-C**). In contrast, neutrophils were the only immune population that accumulated during lactation, increasing from ∼5% to ∼14% of total lung immune cells and exhibiting a 2–3-fold increase in absolute number compared with nulliparous controls (**Fig. 2A, B**).

**Figure 2.**
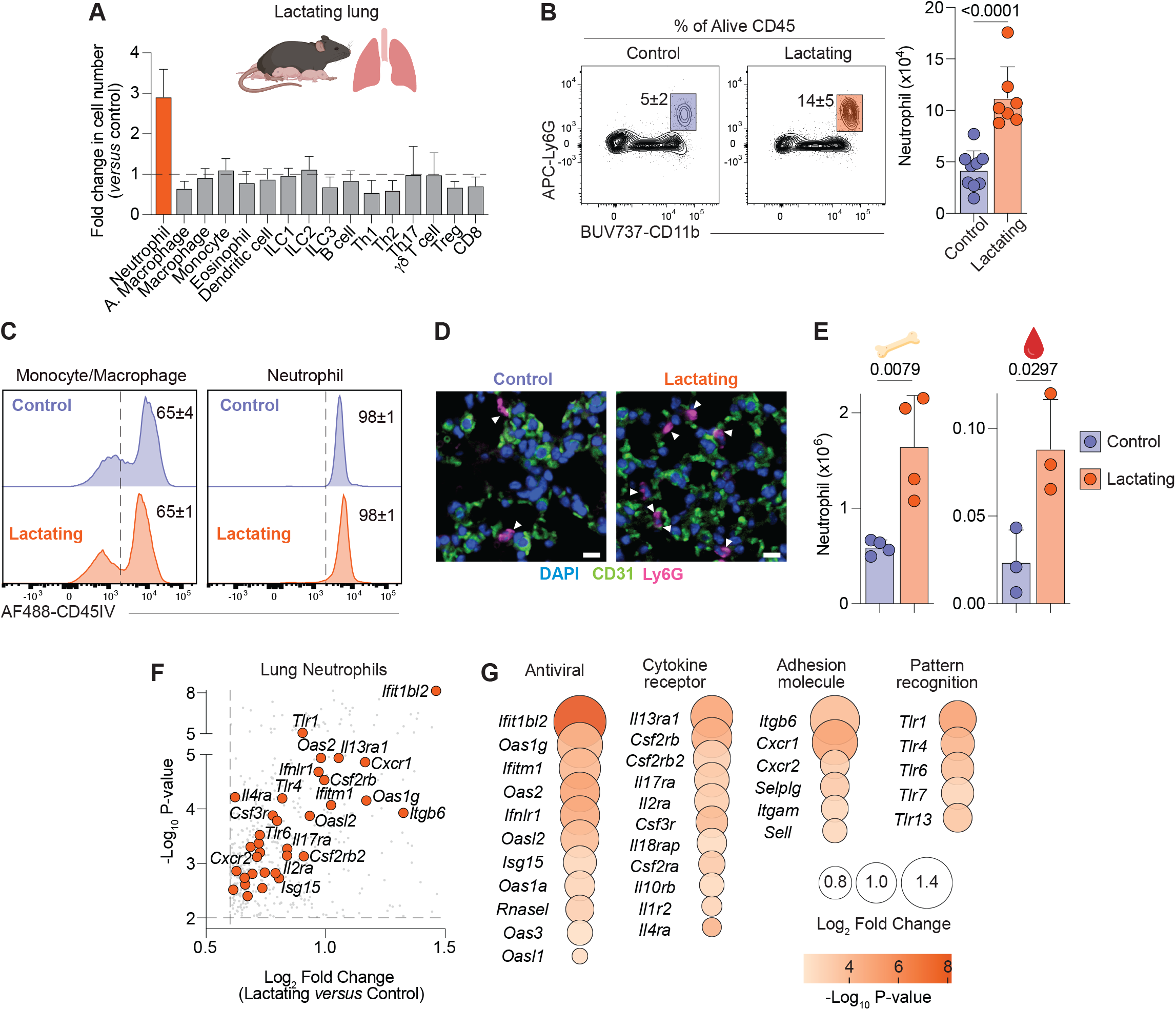
Activated neutrophils accumulate in the lung vasculature during lactation. (**A**) Fold changes of the total cell number of major lung immune cell populations in lactating versus nulliparous mice. (**B**) Representative contour plots (left) of neutrophil frequency (Gated on live CD45^+^CD90.2^-^B220^-^) and absolute cell number (right) in lungs. (**C**) Representative histogram of intravascular CD45 labeling of lung neutrophils. (**D**) Representative immunofluorescence images of lungs. Arrows indicate Ly6G^+^ neutrophils associated with CD31^+^ vasculature. Scale bars, 10 µm. (**E**) Absolute numbers of neutrophils in bone marrow and blood quantified by flow cytometry. (**F**) Volcano plot of differentially overexpressed genes in lung neutrophils from lactating versus nulliparous mice. (**G**) Fold changes and *P* values for defense genes in lung neutrophils from lactating versus nulliparous mice. Data in (B), and (E) represent mean ± SD and were analyzed by unpaired *t*-test. Each dot represents one animal. Data are representative or pooled from two independent experiments. *P* values are indicated.

At homeostasis, pulmonary neutrophils are largely confined to the vasculature and rapidly infiltrate the lung parenchyma and airway upon infection or injury (*26*–*28*). To determine whether lactation-associated neutrophils remained within the pulmonary vasculature or infiltrated the tissue, we performed intravascular immune cell labeling, enabling us to distinguish intravascular from extravascular immune cells. Consistent with previous reports (*29*), neutrophils in nulliparous controls were largely confined to the vasculature, whereas monocytes/macrophages (CD11b^+^ F4/80^+^) were detected in both the vasculature and lung tissues. Despite their marked accumulation during lactation, neutrophils remained predominantly within the vasculature, mirroring their distribution in nulliparous controls (**Fig. 2C**). Thus, lactation is associated with pulmonary intravascular neutrophil accumulation without promoting tissue infiltration. Immunofluorescence imaging further showed increased neutrophil accumulation throughout the lung (**Fig. 2D**).

We next examined whether neutrophil accumulation during lactation was restricted to the lung or extended to other barrier tissues.

Consistent with our previous findings (*13*), neutrophils were modestly increased in the small intestinal lamina propria, whereas neutrophil numbers in the skin remained unchanged (**fig. S3A**). In contrast, neutrophil numbers were significantly increased in both the bone marrow and blood of lactating dams (**Fig. 2E**). Notably, increased bone marrow and lung neutrophils were also observed in *Rag2*^*-/-*^, *Rag2*^*-/-*^*Il2rg*^*-/-*^ and wild-type germ-free lactating mice (**fig. S3B, C**). These findings indicate that enhanced neutrophil production and pulmonary accumulation occur independently of adaptive immunity, innate lymphocytes, and the microbiota.

Building on prior studies demonstrating that neutrophils undergo tissue- and context-specific functional reprogramming (*30*–*32*), we next asked whether their increased number during lactation was accompanied by a distinct transcriptional program. To this end, we isolated neutrophils from the bone marrow and lung of nulliparous and lactating mice and examined their transcriptomes by bulk RNA sequencing (**fig. S4A**). Despite remaining predominantly confined to the pulmonary vasculature, neutrophils isolated from the lactating lung exhibited a transcriptional activation program, including upregulation of antiviral interferon-stimulated genes (*Ifit1bl2, Oas1g, Ifitm1, Oas2*), cytokine receptors (*Csf2rb, Il17ra*), chemokine receptors (*Cxcr1, Cxcr2*), and pattern recognition receptors (*Tlr1, Tlr4, Tlr6*) (**Fig. 2F, G, Table S3**). In contrast, bone marrow neutrophils showed no induction of the activation program observed in the lung, but instead exhibited enrichment of cell-cycle-associated genes (*Sertad4, H3c8, H4c11*) (**fig. S4B, Table S4**). Together, these findings reveal that during lactation, dams harbor enhanced neutrophil production in the bone marrow, accompanied by accumulation of activated intravascular neutrophils in the maternal lung, all in the absence of pathogen exposure.

### Broad protection against respiratory pathogens during lactation

The convergence of epithelial and endothelial remodeling with the accumulation of activated neutrophils led us to hypothesize that lactation establishes a tissue-wide preemptive innate defense program that protects the mother against respiratory infection. To test this, we challenged nulliparous controls and lactating dams with influenza A virus, a major respiratory pathogen (**Fig. 3A**). Consistent with previous reports (*33, 34*), infected nulliparous mice rapidly lost body weight and developed hypothermia beginning two days after infection, hallmarks of progressive disease severity. In striking contrast, lactating dams largely maintained body weight and body temperature throughout the acute phase of infection (**Fig. 3B, C**). This marked improvement in disease outcome was accompanied by significantly lower pulmonary viral burden in lactating dams compared with nulliparous controls, consistent with heightened innate defense established prior to pathogen exposure that limits viral establishment and subsequent expansion (**Fig. 3D**). Strikingly, this protection persisted following lethal challenge, as all nulliparous controls succumbed by day 9 post-infection, whereas 80% of lactating dams survived (**Fig. 3E**).

**Figure 3.**
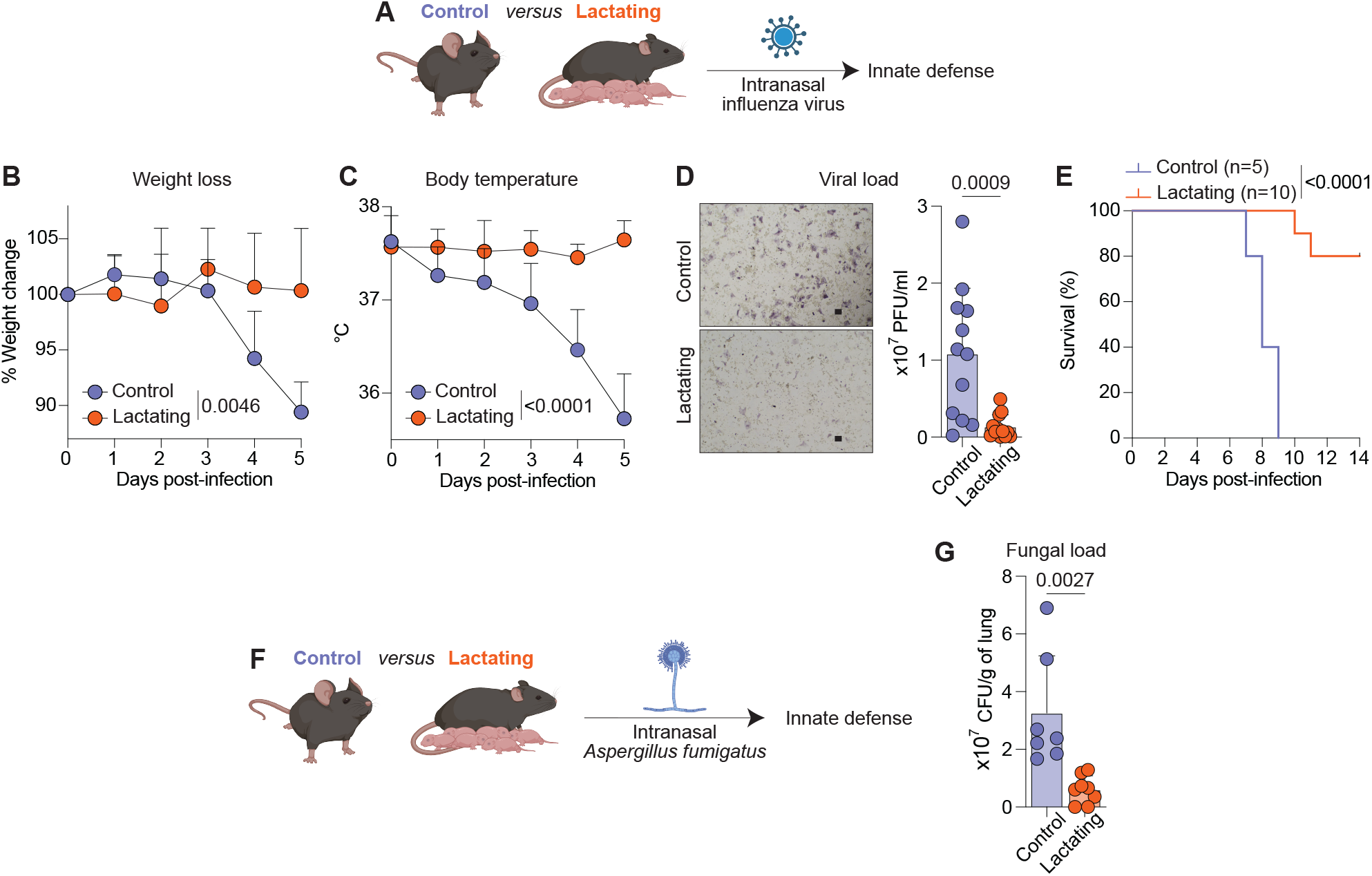
Lactating dams are protected against respiratory viral and fungal pathogens. (**A**) Nulliparous control and day 11 lactating mice were intranasally infected with influenza virus (IAV). (**B**) Body weight changes following sublethal IAV infection (day 0 = 100%) (**C**) Body temperature following sublethal IAV infection. (**D**) Representative image of plaque assay (left) and quantification of viral titer in bronchoalveolar lavage fluid (right) at day 4 post-infection. Scale bars, 100 µm. (**E**) Kaplan-Meier survival curve of nulliparous control and lactating mice infected with 0.5 median lethal dose (LD_50_) IAV. (**F**) Nulliparous control and lactating mice were intranasally infected with the conidia of *Aspergillus fumigatus*. (**G**) Quantification of pulmonary fungal burden at 18 hours post-infection. Data in (B) and (C) were analyzed by paired t-test. Data in (D) and (G) represent mean ± SD and each dot represents one mouse. Data in (D) and (G) were analyzed by unpaired t-test. Data are pooled from at least two independent experiments. Survival analysis in (E) was performed using the log-rank (Mantel-Cox) test. *P* values are indicated.

To determine whether lactation-associated protection extends beyond viral infection, we challenged mice with the conidia of *Aspergillus fumigatus*, an opportunistic fungal pathogen that relies critically on neutrophils and surfactant proteins for clearance (*35, 36*). To assess early pathogen control, we administered a sublethal inoculum and quantified pulmonary fungal burden (**Fig. 3F**). Similar to influenza virus infection, lactating dams exhibited significantly reduced fungal burden compared with nulliparous controls (**Fig. 3G**). Collectively, these findings demonstrate profound protection across viral and fungal respiratory pathogens during lactation, supporting the idea that this tissue-wide preemptive innate defense program is poised to restrict pathogen colonization and enable rapid containment following exposure.

### Prolactin promotes pulmonary neutrophil accumulation during lactation

Having established a functional preemptive innate defense program in the lactating lung, we next sought to determine whether this adaptation is uniquely coupled to active lactation or instead arises following pregnancy and parturition. We therefore compared the epithelial-derived innate defense molecules identified above and neutrophil accumulation across four reproductive conditions, including age-matched nulliparous controls, mid-gestation dams (day 12), peak-lactating dams (day 12), and non-lactating dams. For the non-lactating group, pups were removed within 12–24 hours of birth and mothers were analyzed on day 12 postpartum, allowing dams to experience pregnancy and parturition with minimal lactation (**Fig. 4A**). Strikingly, epithelial-derived surfactant proteins A and D, as well as MUC1, were selectively elevated during active lactation, but not during pregnancy or in non-lactating dams, demonstrating that the epithelial innate defense program is specifically associated with lactation rather than a residual consequence of pregnancy or parturition (**Fig. 4B**).

**Figure 4.**
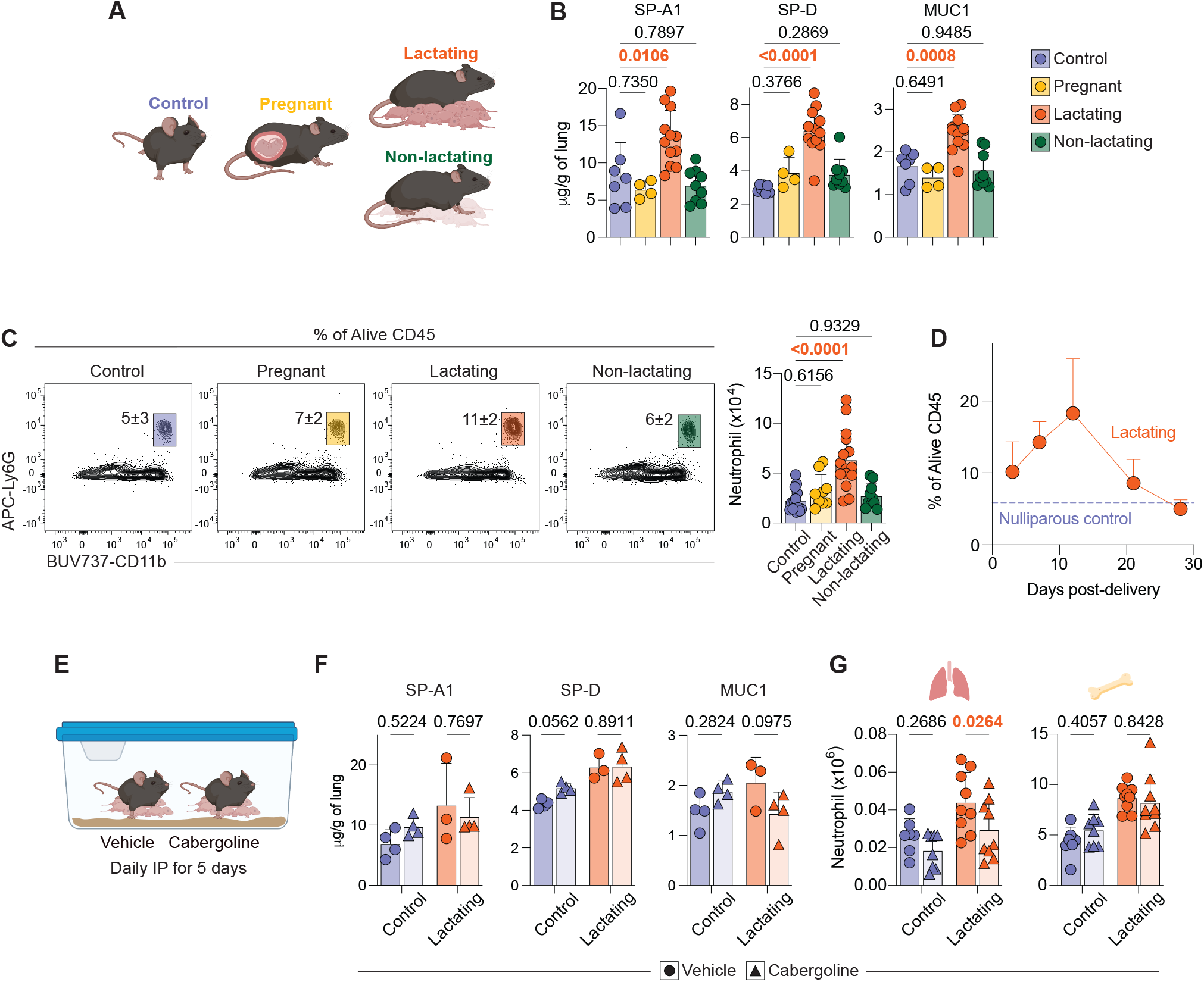
Pulmonary innate defense is selectively established during lactation and regulated in part by prolactin. (**A**) Lungs from nulliparous, day 12 pregnant, day 12 lactating, and day 12 non-lactating mice were analyzed. (**B**) Protein concentrations in lung measured by ELISA. (**C**) Left: Representative contour plots of neutrophil frequency (Gated on live CD45^+^CD90.2^-^B220^-^) and absolute cell number (right) in lungs. (**D**) Neutrophil frequency of total lung immune cells on days 3, 7, 12, 21, and 28 of lactation versus nulliparous control mice. (**E**) Two lactating mice were cohoused and IP-injected with vehicle or cabergoline daily for 5 days. Protein concentrations in lung from vehicle- or cabergoline-treated nulliparous control and lactating mice, measured by ELISA. (**G**) Absolute cell number of lung (left) and bone marrow (right) neutrophils in vehicle- or cabergoline-treated nulliparous control and lactating mice. Data in (B), (C), (F), and (G) represent mean ± SD and each dot represents one mouse. Data in (B) and (C) were analyzed by one-way ANOVA. Data in (F), and were analyzed by two-way ANOVA with Sidak’ s multiple comparisons correction. Data are representative or pooled from two independent experiments. *P* values are labeled.

Similarly, lung neutrophil accumulation was absent in both pregnant and non-lactating dams and occurred exclusively during lactation (**Fig. 4C**). Kinetic analysis further revealed that neutrophil accumulation emerged around day 3 of lactation, progressively increased to peak levels between days 11 and 14, and rapidly declined to levels comparable to those of nulliparous controls by day 21, coinciding with weaning (**Fig. 4D**). Remarkably, the kinetics of neutrophil accumulation closely mirrored the physiological demands of lactation, peaking during maximal breastmilk production and declining thereafter as pups progressively transitioned to solid food. These findings reveal that preemptive innate defense in the maternal lung is a dynamic physiological adaptation whose temporal pattern closely parallels the demands of lactation.

This close temporal relationship led us to hypothesize that prolactin, a principal hormonal regulator of lactation, contributes to pulmonary preemptive innate defense. To test this, we suppressed prolactin secretion with the dopamine receptor D2 agonist, cabergoline, while maintaining pup interactions by co-housing cabergoline-treated dams with vehicle-treated lactating dams (**Fig. 4E**). Consistent with previous reports (*37*), cabergoline treatment resulted in marked mammary gland involution, confirming effective disruption of lactation (**fig. S5A**). Cabergoline treatment had limited effects on epithelial innate defense program, with SP-A and SP-D, and MUC1 remaining elevated (**Fig. 4F**). In striking contrast, lung neutrophil accumulation cabergoline-treated dams, whereas bone marrow neutrophil numbers remained elevated and comparable to those of vehicle-treated lactating dams (**Fig. 4G**). Together, these findings establish enhanced innate defense in the maternal lung as a lactation-specific physiological adaptation, with prolactin signaling selectively regulating pulmonary neutrophil accumulation.

### CGRP contributes to pulmonary remodeling during lactation

We next investigated whether prolactin signaling acts directly within the lung or instead through an intermediary pathway. Analysis of our bulk epithelial and endothelial transcriptomic datasets, together with a published lung single-cell atlas (*38*), revealed little to no detectable expression of the prolactin receptor (*Prlr*) across major epithelial, endothelial, mesenchymal, and immune populations in the lung (**fig. S5B, C**), suggesting that prolactin promotes lung neutrophil accumulation indirectly. Receptors for oxytocin (*Oxtr*) and dopamine (*Drd1a, Drd2, Drd3, Drd4, Drd5*), the latter of which regulates prolactin secretion, similarly exhibited little to no detectable expression across major lung populations (**fig. S5C**), further pointing to alternative pathways contributing to lung neutrophil accumulation during lactation.

To identify candidate signaling pathways that regulate maternal lung remodeling within the intact tissue architecture, we performed Xenium spatial transcriptomic profiling of lungs from nulliparous and lactating dams. Unbiased clustering identified all major epithelial, endothelial, mesenchymal, and immune cell populations, followed by subclustering into their constituent cell population (**Fig. 5A, B, fig. S6A-I**), with their relative abundance broadly consistent with previously published lung cell atlases (*39, 40*). Spatial mapping preserved the anatomical organization of these populations within the lung (**Fig. 5C**). Importantly, the spatial transcriptomic dataset recapitulated our prior flow cytometric analyses, with neutrophils emerging as the only immune population to undergo substantial accumulation during lactation, whereas most other cellular populations remained largely unchanged (**fig. S6J**). These findings suggest that the epithelial and endothelial remodeling observed during lactation is primarily associated with functional reprogramming of existing cellular populations rather than major shifts in cellular abundance or tissue composition.

**Figure 5.**
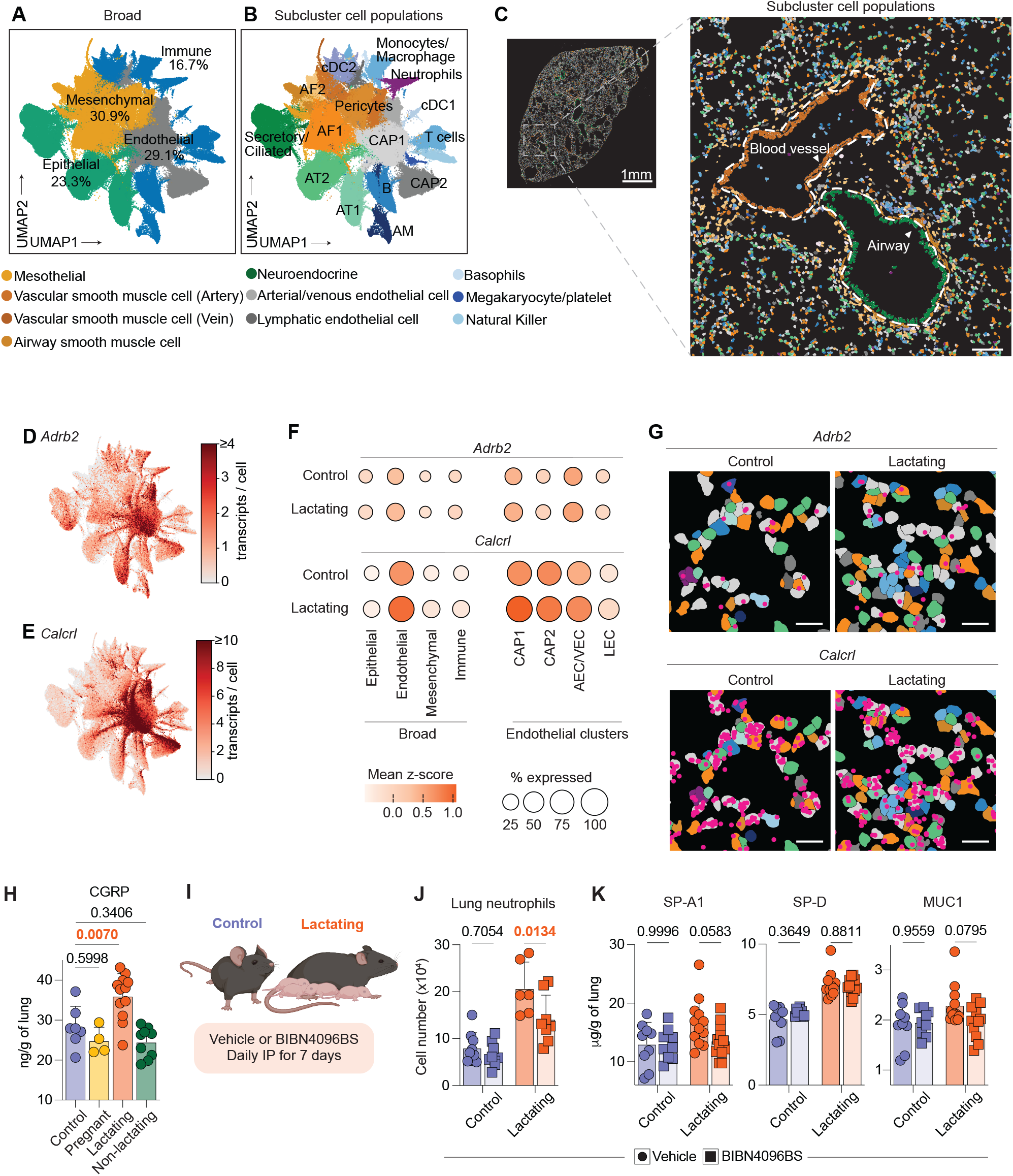
CGRP contributes to pulmonary remodeling during lactation. (**A**) UMAP representation colored by broad cell-type annotation (epithelial, endothelial, mesenchymal, and immune). (**B**) UMAP representation of all profiled cells, colored by Leiden cluster identity. Cell type abbreviations are as follows: AF1, Alveolar fibroblast 1; AF2, Alveolar fibroblast 2; AM, Alveolar macrophage; AT1, Alveolar type 1; AT2, Alveolar type 2; CAP1, Capillary endothelial type 1; CAP2, Capillary endothelial type 2. (**C**) Representative views of Xenium analysis of lung (Control, n = 3; Lactating, n = 3). Scale bars, 100 µm. (**D**) UMAP representation showing *Adrb2* transcripts/cell across Xenium-identified lung cell types. (**E**) UMAP representation showing *Calcrl* transcripts/cell across Xenium-identified lung cell types. (**F**) Percent expressed and mean z-scores for *Adrb2* and *Calcrl* expression across broad cell types and endothelial clusters in lungs of nulliparous and lactating mice. (**G**) Representative visualization of *Adrb2* (top) and *Calcrl* (bottom) expression in the lungs of nulliparous (left) and lactating (right) mice. Scale bars, 20 µm. (**H**) Lung protein concentration of CGRP measured by ELISA. (**I**) Control and lactating mice were IP-injected with BIBN4096BS daily for 7 days. (**J**) Absolute cell number of lung neutrophils in vehicle- or BIBN4096BS-treated nulliparous control and lactating mice. (**K**) Protein concentrations in lung from vehicle- or BIBN4096BS-treated nulliparous control or lactating mice, measured by ELISA. Data in (H), (J), and (K) represent mean ± SD and each dot represents one mouse. Data in (H) were analyzed by one-way ANOVA. Data in (J) and (K), were analyzed by two-way ANOVA with Sidak’ s multiple comparisons correction. Data are pooled from two independent experiments. *P* values are indicated.

We next focused on candidate signaling pathways with established roles in barrier tissue immunity whose receptors showed broad expression across the lung. Among these, *Adrb2*, encoding the β2-adrenergic receptor, and *Calcrl*, encoding the ligand-binding component of the CGRP receptor complex, emerged as potential candidates (**Fig. 5D, E**). Both transcripts were predominantly expressed within the endothelial compartment, although their distribution across endothelial subtypes differed. *Adrb2* was detected in a substantial fraction of CAP1 endothelial cells, as well as arterial and venous endothelial cells (AEC/VEC) while *Calcrl* showed marked expression across endothelial populations (**Fig. 5F, G**).

Building on their established roles in lung immunity (*41*–*46*), we prioritized β-adrenergic and CGRP signaling for functional interrogation. Pharmacological blockade of β-adrenergic signaling with propranolol had minimal effects on lactation-associated lung neutrophil accumulation and bone marrow granulopoiesis, indicating that this pathway is unlikely to be a major driver of enhanced neutrophil production or pulmonary accumulation during lactation (**fig. S7A, B)**. In contrast, pulmonary CGRP levels were selectively elevated during active lactation, but not during pregnancy or in non-lactating dams, closely paralleling the lactation specificity of the preemptive innate defense program (**Fig. 5H**). To determine whether CGRP contributes to lung remodeling during lactation, we treated lactating dams with olcegepant (BIBN4096BS), a CGRP receptor antagonist developed for the treatment of migraine in humans, from postpartum day 5 to 11 (**Fig. 5I**). Despite its lower affinity for the rodent CGRP receptor, olcegepant treatment significantly reduced lung neutrophil accumulation observed during lactation and modestly attenuated the induction of epithelial-derived surfactant protein A and MUC1, whereas surfactant protein D remained unchanged (**Fig. 5J, K**).

These findings reveal that the tissue-wide preemptive innate defense program during lactation integrates multiple regulatory inputs, with prolactin and CGRP contributing predominantly to pulmonary neutrophil accumulation but only partially to the epithelial defense program, pointing to complementary mechanisms that collectively reinforce maternal innate defense during one of the most physiologically demanding periods of maternal life.

### Lactation-associated granulocyte adaptations extend across phylogenetically distant mammals

Together, our findings in mice support a model in which lactation promotes tissue-wide preemptive innate defense in the maternal lung, characterized by enhanced epithelial barrier functions, endothelial remodeling that supports granulocyte recruitment, and accumulation of intravascular neutrophils. This phenomenon is supported by increased bone marrow neutrophil production and systemic mobilization (**Fig. 6A**). To investigate whether this lactation-associated adaptation represents a conserved feature of mammalian reproduction or a species-specific response, we examined circulating granulocytes in phylogenetically distinct mammals. We first sampled the common marmoset (*Callithrix jacchus*), a non-human primate phylogenetically closer to humans than rodents. We reasoned that evidence of a similar response in this lineage would indicate that this adaptation is not restricted to a single mammalian order. In the absence of species-specific immunophenotyping reagents, circulating granulocytes were quantified by flow cytometry based on forward- and side-scatter characteristics (**Fig. 6B**). Remarkably, lactating marmosets harbored significantly increased numbers of circulating granulocytes compared with age-matched nulliparous controls, recapitulating the observation in lactating mice (**Fig. 6B, C**). This suggests that lactation-associated granulocyte expansion extends beyond rodents to primates.

**Figure 6.**
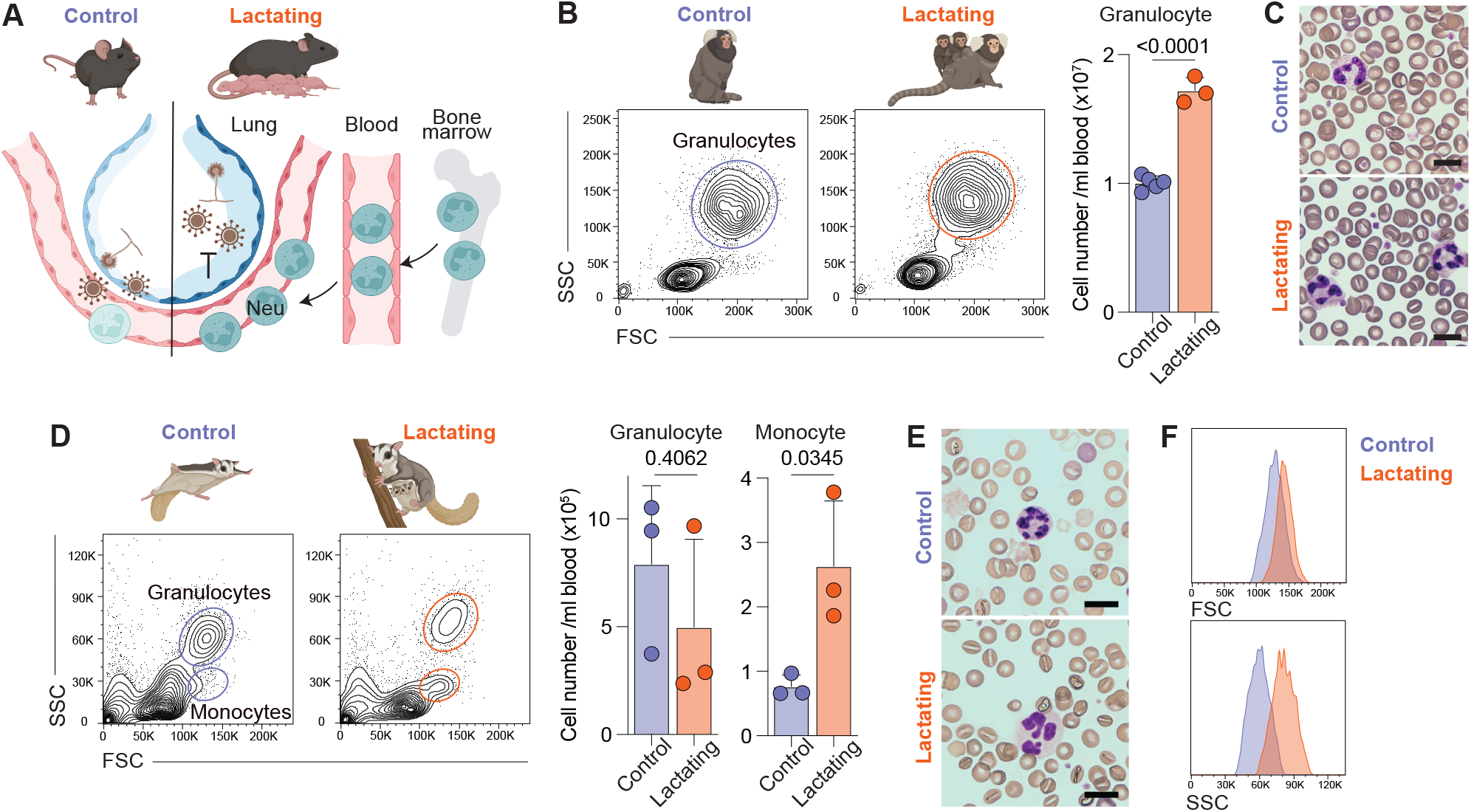
Lactation-associated granulocyte adaptations across mammalian species. (**A**) Model: Lactation promotes tissue-wide preemptive innate defense through increased epithelial-derived surfactants and mucin, endothelial remodeling that supports granulocyte recruitment, and increased bone marrow neutrophil production, systemic mobilization and accumulation of intravascular neutrophils in the lung. Together, these adaptations enhance protection against diverse respiratory pathogens. (**B**) Representative contour plots of circulating granulocytes gated on alive CD45^+^ cells (left) and absolute number of circulating granulocytes quantified by flow cytometry in marmosets (right). (**C**) Representative blood smears of circulating granulocytes in marmosets. (**D**) Representative contour plots of circulating granulocytes and monocytes gated on alive cells (left) and absolute number of circulating granulocytes and monocytes quantified by flow cytometry in sugar gliders (right). (**E**) Representative blood smears of circulating granulocytes in sugar gliders. (**F**) Representative histogram of FSC and SSC of circulating granulocytes in sugar gliders. Data in (B), and (D) represent mean ± SD and were analyzed by unpaired *t*-test. Each dot represents one animal. *P* values are indicated.

Having established that lactation-associated granulocyte expansion extends to a non-human primate, we next asked whether this phenomenon represents a broader, ancestral feature of mammalian reproductive biology. We therefore turned to the sugar glider (*Petaurus breviceps*), a marsupial that diverged from eutherian mammals more than 160 million years ago (*47*–*49*). Because marsupials represent an early-diverging mammalian lineage, examining this species allowed us to ask whether lactation-associated granulocyte adaptations might predate the eutherian-marsupial split, rather than having arisen independently within eutherian mammals. In sugar gliders, circulating granulocyte numbers remained unchanged during lactation, although monocyte numbers were increased (**Fig. 6D**). Despite the absence of numerical expansion, blood smear and flow cytometric forward- and side-scatter analyses revealed that lactating sugar gliders harbored circulating granulocytes with increased size and granularity (**Fig. 6E, F**), a morphological characteristic associated with activation (*50, 51*). Thus, although the granulocyte adaptation differed across species, with numerical expansion in mice and marmosets and activation-associated morphological changes in sugar gliders, all three phylogenetically distinct mammals exhibited lactation-associated granulocyte adaptations. These findings raise the possibility that granulocyte adaptations represent an evolutionarily conserved feature of lactation.

## Discussion

Lactation is a defining feature of mammals and one of the most physiologically demanding periods of maternal life. Here, we show that lactation equips the maternal lung with a preemptive innate defense program and protects mothers against respiratory viral and fungal pathogens. Rather than relying largely on responses mounted after pathogen encounter, the lactating lung is remodeled in advance across epithelial, vascular, and immune compartments for a multilayered defense program poised to limit pathogen entry and enable rapid containment following exposure.

Along with our previous discovery of eosinophil-driven maternal intestinal remodeling (*13*), these findings demonstrate that in parallel to systemic immune modulation during reproduction, maternal barrier tissues mount distinct local defense programs. In the intestine, eosinophils accumulate starting in pregnancy, peak during lactation, and persist long-term postpartum, promoting goblet cell differentiation and mucosal defense. In contrast, pulmonary remodeling is restricted to active lactation, with heightened epithelial production of surfactant proteins and mucin, along with prominent neutrophil accumulation, which rapidly declines with weaning. Thus, even within the same reproductive cycle, individual barrier organs operate on distinct temporal programs and employ different cellular strategies to enhance maternal immune defense. Notably, both programs engage granulocytes yet deploy distinct populations to support tissue-specific functions, revealing a division of labor across maternal barrier tissues. Why pulmonary remodeling selectively occurs during lactation, whereas intestinal remodeling begins during pregnancy and persists through the late postpartum period, remains unknown. These divergent kinetics suggest that individual barrier tissues respond to distinct physiological cues, allowing maternal barrier immunity to be dynamically tailored throughout the reproductive cycle.

Our findings further identify a role for neuroendocrine signaling in lactation-associated lung remodeling. Prolactin and CGRP signaling contributed prominently to pulmonary neutrophil accumulation while exerting modest effects on epithelial remodeling. The cellular source of CGRP during lactation remains to be defined and may include rare neuroendocrine populations, which together with sensory neurons, are established sources of CGRP in response to airway stimuli (*52*–*54*). Notably, the differential and incomplete effects of prolactin and CGRP point to a distributed regulatory system in which complementary signals collectively establish tissue-wide protection. The additional signals shaping the broader epithelial defense program remain unknown and may involve other neuroendocrine, metabolic, or tissue-derived cues associated with lactation. Identifying these additional regulatory inputs will be important for understanding how the physiological state of lactation is translated into tissue-specific immune adaptation and may reveal broader principles of tissue immunophysiology during reproduction.

Intriguingly, the lactation-associated granulocyte adaptations were observed across marsupials and eutherian mammals, two lineages that diverged more than 160 million years ago. This broad evolutionary distribution raises the possibility that granulocyte adaptation is a deeply conserved feature of mammalian reproductive biology. Although this phenomenon differs across species, our findings point to a shared evolutionary strategy in which lactation is accompanied by granulocyte adaptations, highlighting maternal innate defense as an integral component of mammalian reproductive biology. In natural environments, where mammals evolved under persistent pathogen pressure, such granulocyte adaptations may have conferred an evolutionary advantage by strengthening maternal innate defense and thereby promoting survival of both mother and the offspring.

Together, our work broadens the framework of reproductive immunology beyond systemic immune modulation that supports fetal tolerance, revealing tissue-specific immune adaptations that safeguard maternal health against pathogen exposure. Our findings further demonstrate that enhanced maternal innate defense emerges from integrated adaptations across cell types and regulatory networks that collectively meet the changing physiological demands of reproduction. In light of these observations, our work highlights the need for a deeper understanding of how distinct physiological states in reproduction shape maternal tissue function, disease susceptibility, and therapeutic responses, knowledge that is essential for advancing maternal health throughout reproduction.

## Acknowledgments

We thank all members of Lim laboratory for insightful discussion and constructive feedback; staff at Princeton University and Howard Hughes Medical Institute for their continuous assistance. We thank staff at the Princeton Molecular Biology flow cytometry resource facility, confocal imaging facility, the Lewis-Sigler Institute Genomics core facility, and Laboratory Animal Resources for technical support. Flow data were acquired using the BD Biosciences FACSymphony A3 Flow Cytometer, purchased with an NIH S10 award (1S10OD028592-01A1). We thank Michel Enamorado for valuable scientific discussion and advice.

## Author Contributions

Conceptualization: A.I.L., T.R.C. Investigation: T.R.C., C.Y.H., S.L., K.H.A.F., R.S., D.S., H.S., J.R.d.S., L.Y., Methodology: A.I.L., T.R.C., S.L., K.H.A.F. Supervision: A.I.L., J.V.D., F.M.K., R.M. Visualization: A.I.L., T.R.C., S.L. Writing-original draft: A.I.L., T.R.C. Writing-reviewing and editing: all authors and all Lim Laboratory members.

## Materials and methods

### Mice

C57BL/6J (strain no. 000664) and *Rag2*^*-/-*^ (strain no. 008449) mice were purchased from Jackson Laboratory. *Rag2*^*-/-*^*Il2rg*^*-/-*^ (model no. 4111) and Germ-free (GF) mice (model no. B6) were purchased from Taconic Bioscience. All experiments described in this study were performed in accordance with protocols that were approved by the Institutional Animal Care and Use Committee (IACUC) of Princeton University under protocol number 3087. All mice were bred and maintained in an AAALAC-accredited animal facility at Princeton University. Tissue collection and analysis was performed on gestation day E12.5 for pregnant mice or 12 days postpartum for lactating and non-lactating mice. Age-matched mice that were co-housed with males but did not get pregnant were used as nulliparous controls.

### Sugar gliders

Experiments were performed in accordance with protocols approved by the IACUC of Princeton University under the protocol number 2155. Adult captive-born sugar gliders were acquired from the US pet trade and housed in a breeding colony at Princeton University. Animals were housed and maintained as previously described (*55*). Briefly, animals were fed a diet consisting of dried food, fruits, and protein daily, and were housed in bonded pairs or trios. The colony was on a 12 hr light-dark cycle with a room temperature of 20-27°C and humidity of 30-70%.

### Marmosets

Common marmosets were housed in family groups under controlled environmental conditions with a room temperature maintained between 24-30°C, relative humidity between 30-70%, and a 12-hour light-dark cycle in accordance with protocols approved by the IACUC of Princeton University under protocol number 3118.

### Cell line

Madin-Darby canine kidney (MDCK, CCL-34, ATCC) cells were grown in Dulbecco’ s Modified Eagle Medium (DMEM) supplemented with 10% fetal bovine serum (FBS), 2mM L-glutamine, 100 U/ml penicillin, and 100 µg/ml streptomycin at 37ºC and 5% CO2. Cells were passaged every 2-3 days using 0.05% trypsin-EDTA. MDCK cells were used for influenza virus experiments described below.

### Influenza infection

Control or day 11 lactating mice were infected with Influenza A virus PR8 (VR-1469, ATCC). Mice were anesthetized with isoflurane and intranasally infected with 0.1 (sublethal) or 0.5 (lethal) LD50 PR8 in 40 µl sterile 1X Phosphate-Buffered Saline (PBS). For monitoring experiments, mice were monitored for weight, temperature, and clinical symptoms for 5 days. For survival experiments, mice were monitored for 14 days and euthanized if weight loss exceeded 25%. IAV viral titers from bronchoalveolar lavage fluid (BALF) were quantified using an immunostaining-based plaque assay as previously described (*56*). Briefly, 1 ml of 1X PBS was used to flush the airways of infected mice and BALF was collected and centrifuged at 500 x g for 5 min at 4ºC. MDCK cells were plated in a 24-well plate in DMEM supplemented with 10% FBS. BALF supernatant was serially diluted in serum-free DMEM, and 200 µl was used to inoculate the cells for 1 hr at 37ºC and 5% CO2. Cells were then overlayed with 500 µl of DMEM/F12 (12-500-062, Gibco) containing 1% carboxymethylcellulose (C5678, Millipore-Sigma) and 1µg/ml TPCK-treated trypsin (20233, Thermo Fisher) and incubated for 72 hr at 37ºC and 5% CO2. Cells were then fixed with ice-cold methanol for 1 hr and stained with rabbit anti-IAV antibody (1:500, PA5-32242, Invitrogen), followed by staining with HRP-conjugated goat anti-rabbit secondary antibody (1:1000, A16096, Thermo Fisher). Plaques were visualized using HRP substrate and were manually counted under a light microscope. Viral titers were expressed as PFU/ml of BALF.

#### *A. fumigatus* infection

*Aspergillus fumigatus* strain B-5233 conidia were grown and harvested as previously described (*57*). Briefly, conidia were cultured on glucose minimal medium agar slants containing trace elements and salts (*58*) for 5-7 days at 37°C. Subsequently, conidia were harvested by washing the slants with phosphate-buffered saline (PB500, Alkali Scientific) containing 0.01% Tween-20 (P9416, Millipore-Sigma). The resulting conidial suspension was passed through a 70-μm nylon cell strainer (MT4070, Alkali Scientific), washed twice with 0.01% Tween-20 in PBS, counted using hemocytometer and resuspended at a desired concentration for downstream application. Control or day 11 lactating mice were anesthetized with isoflurane and intranasally infected with 5 × 10^7^ conidia in 40 µl sterile 1X PBS with 0.05% Tween-20. Lungs were collected at 18 hr post-infection for CFU quantification. Mouse lungs were weighed and collected in 1ml 1X PBS and homogenized with a Precellys 24 homogenizer for two cycles of 5,000 rpm with a 10 sec break in between. Homogenized lung tissue was serially diluted and plated as triplicates on Malt Extract Agar (70145, Sigma-Aldrich). Plates were incubated for 20 hr at 37ºC and quantified.

### Tissue processing

Mice were euthanized by CO2 euthanasia and tissues (lung, small intestine, skin, and bone marrow) were collected and stored in complete media (RPMI 1640 containing 20 mM HEPES, 2 mM L-glutamine, 1 mM sodium pyruvate, 1mM nonessential amino acids, 50mM β-mercaptoethanol, 100 U/ml penicillin, 100 mg/ml streptomycin, and 3% FBS until processing. For the lung, whole lung was minced and digested in RPMI 1640 supplemented with 1 mg/ml collagenase IV (C5148, Millipore-Sigma), and 0.5 mg/ml DNAse I (DN25, Millipore-Sigma) for 35 min with gentle agitation at 37°C. Remaining cells were then passed through a 70 µm strainer, washed with complete media, and prepared for flow cytometry. For the small intestine lamina propria (siLP), Peyer’ s patches were removed and the tissue was opened and washed with cold 1X PBS to remove fecal contents. Tissue was cut into 1 cm segments and placed into complete media containing 5 mM EDTA and 0.145 mg/ml dithiothreitol (DTT) for 20 min at 37°C with continuous stirring. Next, tissue was vigorously shaken for 1 min in RPMI containing 20 mM HEPES, 100 U/ml penicillin and 100 mg/ml streptomycin for 1 min and further digested with 10 ml of media containing 0.5 mg/ml DNAse I and 0.1 mg/ml Liberase TL (5401020001, Millipore-Sigma) for 25 min at 37°C with constant stirring. Cells were passed through a 70 µm strainer and leukocytes were enriched using 37.5% Percoll. Following a wash with complete media, cells were prepared for flow cytometry. For the skin, ear pinnae were excised and separated into ventral and dorsal sheets, digested by placing the dermal side down in medium containing 500 mg/ml DNase I with 250 mg/ml Liberase TL, and incubated for 90 min at 37°C. Digested ears were homogenized, filtered through a 70 µm strainer, washed in complete media, and prepared for flow cytometry.

### Blood collection and processing

All blood for flow cytometry and blood smear analysis was collected in an EDTA tube and mixed well. For mice, control or day 12 lactating animals were restrained, and the submandibular vein was punctured with a 5mm lancet. For sugar gliders, control or 4–8-week postpartum lactating animals were anesthetized with isoflurane, and a 5mm lancet was used to puncture the skin above the saphenous or cephalic vein. For marmosets, control or ∼4-week postpartum lactating animals were manually restrained and bled from the right or left femoral vein.

### Lung epithelial and endothelial cell isolation

Mice were euthanized by CO2 euthanasia, and the heart was perfused with 10 ml cold Hank’ s Balanced Salt Solution (HBSS) and tied off with suture thread to prevent leakage. The trachea was catheterized and washed three times with 1 ml 0.5M EDTA in 1X PBS. 1 ml digestion solution (DMEM/F12 containing 5 U/ml Dispase II (17105041, Gibco), 4.5 U/ml Elastase (E7885, Millipore-Sigma), 1mg/ml collagenase IV, and 0.1 mg/ml DNase I) was added, quickly followed by 0.5 ml warm 1% agarose, and the trachea was tied off to prevent leakage. The heart-lung block was placed in digestion solution on ice until further processing. Lung was incubated for 40 min at 37ºC with gentle agitation. Then, lung was separated from the heart and connective tissue, transferred into fresh digestion solution and minced and incubated for 15 min at 37 ºC with gentle agitation. 4 ml of 10% FBS in RPMI was added to stop digestion and incubated on ice for 5 min. The cell solution was then filtered through a 70 µm cell strainer and incubated for 5 min at 37ºC at 100rpm. Cells were then centrifuged for 10 min at 4ºC at 300 x g. Supernatant was aspirated and the pellet was resuspended in 1 ml of ACK red blood cell lysis buffer. The remaining cells were washed with 9 ml 5% FBS in 1X PBS and centrifuged for 10 min at 4ºC at 300 x g and prepared for cell sorting.

### Intravenous immune cell labeling

Intravascular (IV) immune cell labeling was carried out as previously described (*29*). Briefly, nulliparous control and lactating mice were anesthetized and IV injected with 150 µl of anti-CD45.2 antibody and euthanized after 3 min. Lungs were harvested and prepared for flow cytometry.

### Flow cytometry analysis

Cell suspensions were added to a 96-well plate with counting beads. Cells were washed twice in HBSS and stained with primary antibodies for 20 min on ice. Cells were then washed and stained with Live/Dead (1:400, L23105, Invitrogen) for 10 min on ice, followed by another HBSS wash, and 30 min of fixation (00-5123-43, eBioscience) at room temperature. Cells were then washed twice with 1X PBS and analyzed using a BD FACSymphony A3. All antibodies used in this study are listed in **Table S5**.

### Cell sorting

Lung epithelial and endothelial cell, and bone marrow and lung neutrophil suspensions were prepared as described above. All cell suspensions were stained with 4’, 6-diamindino-2-phenylindole (DAPI) (1:200, D3571, Invitrogen) immediately before sorting. Lung epithelial cells were sorted as alive CD45^-^EpCAM^+^CD31^-^. Lung endothelial cells were sorted as alive CD45^-^ EpCAM^-^CD31^+^. Bone marrow and lung neutrophils were sorted as alive CD45^+^CD11b^+^Ly6G^+^. Cells were sorted using a SONY MA900 sorter directly into cell lysis buffer.

### Immunofluorescence imaging of lung

Lungs were fixed in 10% neutral-buffered formalin for 24 hr at 4ºC and embedded in paraffin. 5 µm sections were mounted onto glass slides and deparaffinized and rehydrated. For antigen retrieval, slides were placed into a jar containing target retrieval solution (GV80511-2, Agilent) and placed into a pressure cooker for 40 min at 97 ºC. Slides were then cooled to room temperature, washed with 1X PBS, and permeabilized for 10 min at room temperature with 0.1% Triton X-100 (X100, Millipore-Sigma) in 1X PBS. Next, slides were blocked for 30 min at room temperature in 1X PBS containing Tween-20 and 3% BSA (A3059, Millipore-Sigma), followed by overnight primary antibody staining at 4ºC. The following day, slides were washed three times with 1X PBS for 15 min, stained with DAPI, mounted with Prolong Gold Antifade reagent (P36930, Invitrogen) and dried overnight before imaging on a Nikon CSU-W1.

### Mammary gland imaging

Mammary glands from nulliparous control and lactating mice were gently excised using a scalpel and spread onto a glass slide. Tissue was dried for 1 min at room temperature and fixed overnight in 10% neutral-buffered formalin. Tissue was rehydrated in 75%, 35%, and 15% ethanol, followed by water for 15 min each. Next, mammary gland was stained in carmine solution (07070, Stemcell) overnight at room temperature, followed by dehydration in 70%, 95%, and 100% ethanol for 15 min each. Stained slides were then stored in Histo-Clear (5089990147, Fisher Scientific) until imaging on the Leica Aperio VERSA with Zen software.

### Prolactin blocking treatment

The dopamine D2 receptor agonist, cabergoline (C0246, Sigma-Aldrich), was dissolved in dimethyl sulfoxide (DMSO, BP231, Thermo Fisher) to 10 mg/ml and further diluted in sterile 1X PBS for injection. Nulliparous control and day 7 lactating mice were injected intraperitoneally (IP) with 5 mg/kg cabergoline in 100 µl daily for 5 days. Mice were sacrificed and tissues were analyzed 24 hr after the final dose.

### CGRP blocking treatment

The CGRP receptor antagonist, BIBN4096BS (SML2426, Sigma-Aldrich) was dissolved in DMSO to 2 mg/ml and further diluted in sterile 1X PBS for injection. Nulliparous control and day 5 lactating mice were injected IP with 300 µg/kg BIBN4096BS in 100 µl daily for 7 days. Mice were sacrificed and tissues were analyzed 24 hr after the final dose.

### β-adrenergic receptor blockade

The β1/β2 adrenergic receptor antagonist, propranolol (P0884, Sigma-Aldrich) was diluted in water to 50 mg/ml and further diluted in drinking water. Nulliparous control and day 5 lactating mice were maintained on drinking water containing 0.5 mg/ml propranolol for 7 days. Mice were sacrificed and tissues were analyzed on the final day of treatment.

### Blood smear

5 µl of EDTA-treated whole blood was pipetted onto a glass slide. A second slide held at a 45º angle was backed into the blood drop and streaked across the slide. Slides were dried at room temperature, fixed, and stained with HemaDiff Rapid Stain (Fisher Scientific, STL1197016).

Slides were then washed with water, dried at room temperature, and mounted with Permount (SP15-100, Fisher Scientific). Blood smear images were acquired on a Nikon Eclipse TI-E equipped with a Nikon DsRi2 color camera.

### Bulk RNAseq

Lung epithelial and endothelial cells, and bone marrow and lung neutrophils were isolated from the lung and sorted as described above. Total RNA was extracted from cells using the RNeasy Micro Kit (74034, Qiagen) and analyzed using a Qubit fluorometer and an Agilent Bioanalyzer. Samples with an RNA integrity number (RIN) >7 were submitted to Novogene for analysis. Libraries were prepared using the NEBNext® Ultra™ II RNA Library Prep Kit (New England Biolabs) and were quantified, pooled, and sequenced on the Illumina NovaSeq 6000 platform using a paired-end 150 bp configuration. Sequencing reads were aligned to the *Mus musculus* reference genome (GRCm38/mm10), and transcript abundance was performed by Novogene using standard pipelines. Differential gene expression was determined using DESeq2 and Gene ontology (GO) enrichment was performed using clusterProfiler.

### Spatial transcriptomic analysis of lung using Xenium Prime Mouse 5K

Lung was inflated using a 50% OCT/PBS solution. The inferior lobe was cut in half and embedded in OCT. Tissues were cryosectioned at 10 μm, mounted on Xenium slides and processed according to the Xenium Prime 5K protocol (10X Genomics). Xenium datasets were generated using a Xenium Analyzer (instrument software v.3.4.1.0 and analysis v.xenium-3.3.0.1). Tissue slides were also stained with Xenium MultiTissue Stain mix for multimodal cell segmentation, and this segmentation from Xenium Analyzer was used for all analyses.

Xenium output files were loaded using spatialdata-io (v.0.5.1) and spatialdata (v.0.6.1). Transcripts for each cell were filtered to retain only those overlapping the nucleus segmentation. Cells from all samples were concatenated and filtered to retain cells expressed at least ten genes with a segmented nucleus. Genes detected in fewer than ten cells were also excluded. Data were processed using Scanpy (v.1.11.5). Counts were normalized to library size, log-transformed and then z-scored. Normalized data were used for principal component analysis (PCA) (n_pcs=50 components), *k*-nearest neighbor graph construction (*k*=15) and Leiden clustering (resolution = 0.50), with UMAP embeddings computed using 200 iterations. A cluster with low transcript counts was excluded and clustering was repeated on the remaining cells. Marker genes were identified per cluster by Wilcoxon rank-sum test (one-versus-rest) using ‘ rank_genes_groups’ function, with counts normalized to 10,000 and log transformed. *P* values were corrected with the Benjamini– Hochberg procedure.

Clusters were annotated manually using known marker genes cross-referenced with the LungMAP single-cell reference (CellRef) (*38*) and assigned to four broad cell-type populations (epithelial, endothelial, mesenchymal and immune). Clusters that could not be confidently assigned to any of these populations were excluded for subsequent analysis. Each population was independently reclustered using PCA (50 components), *k*-nearest neighbor graph construction (*k*= 15; n_pcs = 10 for epithelial and endothelial cells, 15 for mesenchymal and immune cells), followed by Leiden clustering at cell-type-specific resolutions (epithelial and endothelial: 0.20; mesenchymal: 0.25; immune: 0.50), with UMAP embeddings computed using 200 iterations. Marker genes were identified per sub-cluster by Wilcoxon rank-sum test (one-versus-rest) as described above. Sub-clusters were manually annotated using known marker genes and CellRef. The top two marker genes per cluster were visualized as dot plots showing mean scaled expression and the fraction of cells expressing each gene. Cell frequencies per cluster were calculated as the total number of cells assigned to each cluster across all samples. To assess changes in cell-type abundance between conditions, cell density (cells per µm^2^) was calculated by dividing cluster cell counts by the tissue section area, manually delineated in Xenium Explorer (10x Genomics). Fold change between lactating and nulliparous conditions was computed as the ratio of mean densities.

### Visualization of *Adrb2* and *Calcrl* expression

Expression of *Adrb2* and *Calcrl* was visualized on the UMAP embedding of all cells. Cells were colored by the number of nucleus-overlapping transcripts detected per cell, with the color scale capped at approximately the 99th percentile for each gene. For each broad population and endothelial subtype, the percentage of cells with at least one transcript and the mean *z*-scored expression were calculated for each condition, pooling cells from all sections of that condition. Dot plots were generated in R (v.4.5.0) using ggplot2 (v.4.0.3).

### Analysis of the LungMAP reference

Expression of lineage marker genes (*Epcam, Pecam1, Ptprc* and *Col1a1*), *Prlr, Oxtr* and dopamine receptor genes (*Drd1a, Drd2*–*Drd5*) was examined in the LungMAP mouse lung single-cell reference (CellRef v.1.1) (*38*). Cells from the endothelial, epithelial, immune and mesenchymal lineages were displayed on the UMAP embedding provided with the reference and colored by the log-normalized expression values supplied with the dataset, using a common color scale for all genes.

### Reference single-cell dataset visualization

To examine the expression of prolactin receptor (*Prlr*), oxytocin receptor (*Oxtr*), and dopamine receptor (*Drd1a, Drd2, Drd3, Drd4, Drd5*) genes across major lung cell types, we used the publicly available CellRef (v1.1) comprising 71,439 cells. Cells annotated as epithelial, endothelial, mesenchymal, and immune were retained for visualization. Pre-computed UMAP coordinates provided in the reference object were used directly. Log-normalized gene expression values were projected onto the UMAP for each gene of interest, together with established broad cell-type marker genes (*Epcam*, epithelial; *Pecam1*, endothelial, *Ptprc*, immune; *Col1a1*, mesenchymal). Visualization was performed using Python (v3.x) with scanpy (v 1.11.5) and matplotlib (v3.10.7).

### ELISA and Luminex

Mouse lung was weighed and collected in 1 ml 1X protease inhibitor (78430, Thermo Fisher) in RIPA buffer (R0278, Sigma-Aldrich) and homogenized for two cycles of 5,000 rpm with a 10 sec break in between. Samples were spun down at 16,000 x g for 10 min at 4ºC and the supernatant was aliquoted and stored at -80 ºC before use. SFTPA1 concentrations were measured using a Mouse Surfactant Protein A ELISA Kit (NBP2-76693, Novus Biologicals), SFTPD was measured using a Mouse SP-D Quantikine ELISA Kit (MSFPD0, R&D systems), MUC1 was measured using a Mouse MUC1 ELISA kit (NBP2-76697, Novus Biologicals), and CGRP was measured using a Rat CGRP1 ELISA Kit (E-EL-R0135, Elabscience). CXCL2 and G-CSF concentrations were measured using a custom Luminex Discovery Assay (R&D systems).

### Statistical analysis

All statistical analyses were performed using GraphPad Prism software. Specific tests used are listed in the figure legends.

## Supplementary Figures

**Figure S1.**
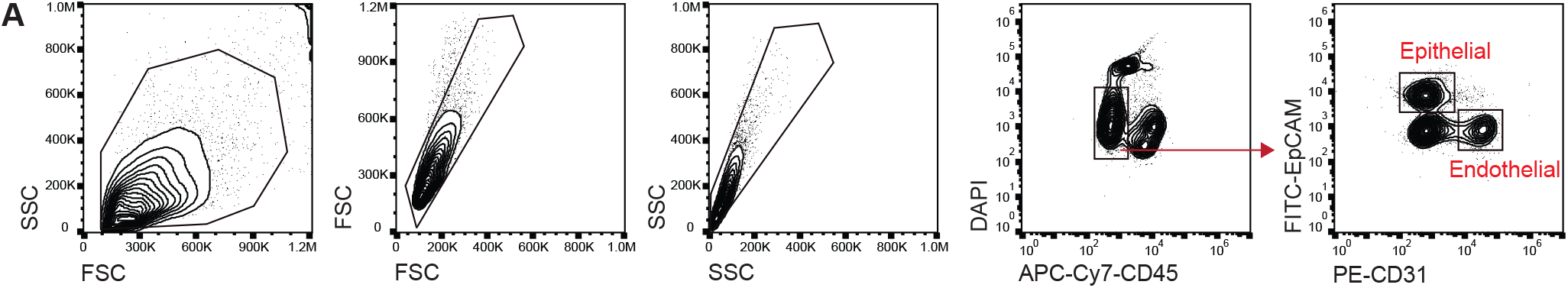
Gating strategy for lung epithelial and endothelial cell sorting. (**A**) Representative flow cytometry gating strategy used to sort lung epithelial and endothelial cells.

**Figure S2.**
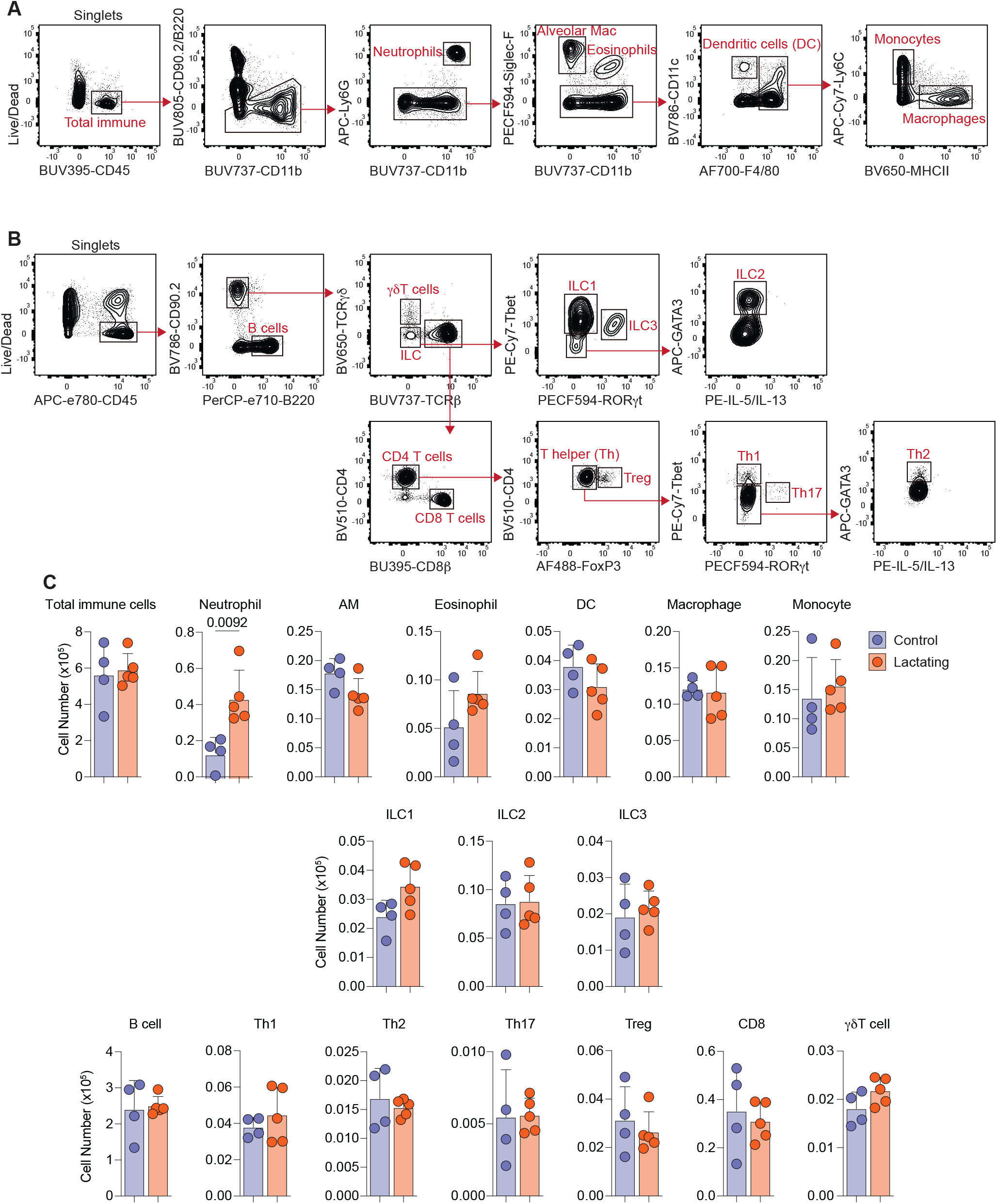
Lactating mice selectively increases neutrophils in the lung. (**A**) Flow cytometry gating scheme for myeloid cells. (**B**) Flow cytometry gating scheme for lymphoid cells. (**C**) Absolute cell number of myeloid and lymphoid cells in the lung of nulliparous control and lactating mice. Data in (C) represents mean ± SD and each dot represents one mouse. Data in (C) were analyzed by unpaired t-test. *P* values are indicated.

**Figure S3.**
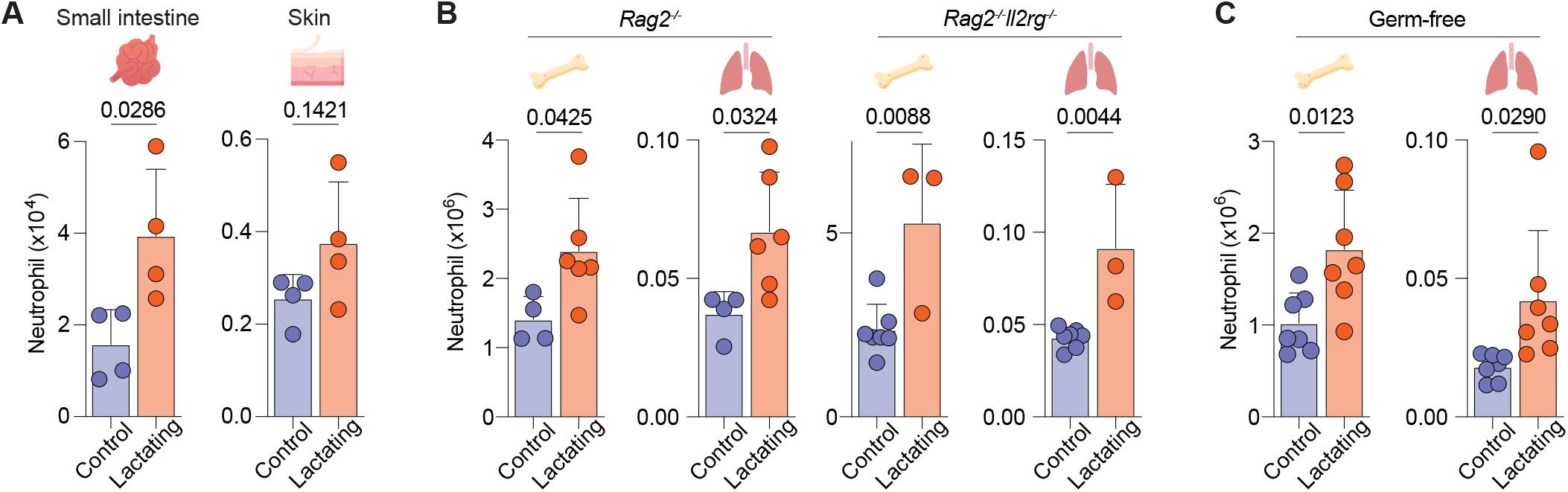
Lactation-associated granulocyte adaptations across tissues and experimental models. (**A**) Absolute cell number of neutrophils in small intestine and skin of nulliparous control and lactating mice. (**B**) Absolute cell number of neutrophils in bone marrow and lung of nulliparous control and lactating *Rag2*^*-/-*^ (left) and *Rag2*^*-/-*^*IL2rg*^*-/-*^ (right) mice. (**C**) Absolute cell number of neutrophils in bone marrow and lung of germ-free nulliparous control and lactating mice. Data in (A), (B), and (C) represent mean ± SD and were analyzed by unpaired t-test. Each dot represents one animal. Data are representative of at least two experiments. *P* values are indicated.

**Figure S4.**
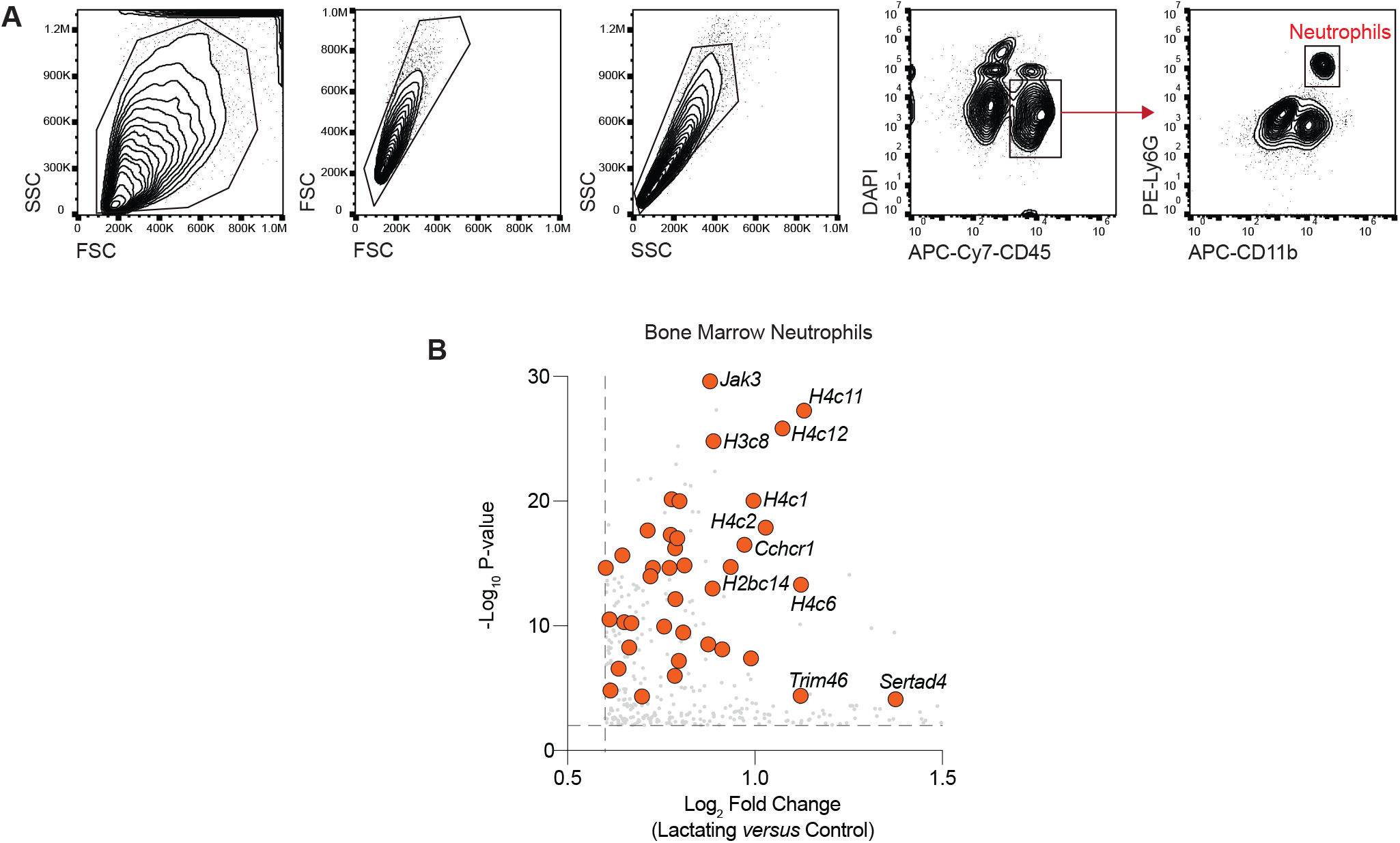
Neutrophils acquire activated signature following bone marrow egress. (**A**) Gating scheme of lung and bone marrow neutrophils. (**B**) Volcano plot of differentially overexpressed genes in bone marrow neutrophils from lactating versus nulliparous mice.

**Figure S5.**
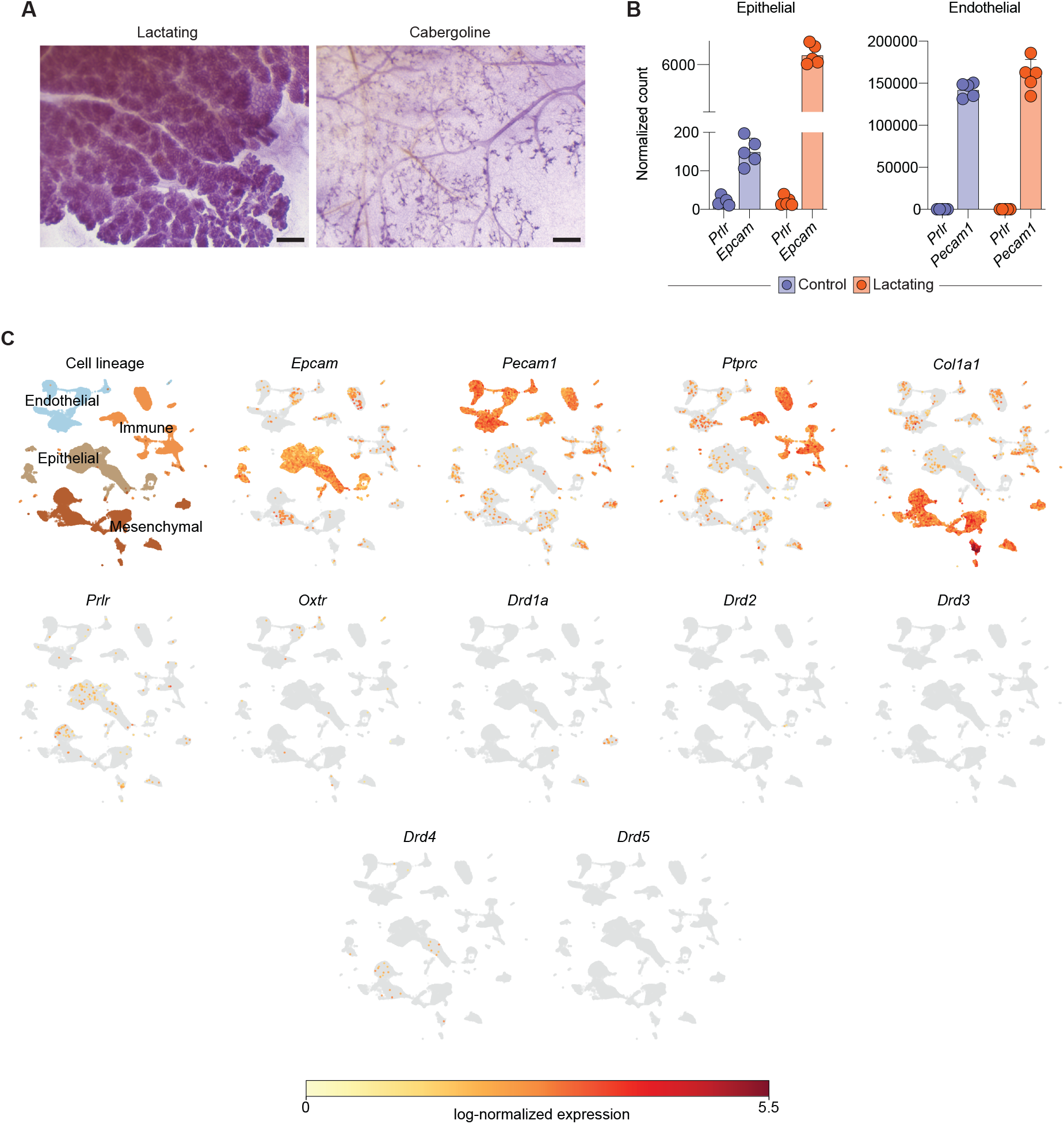
Prolactin, oxytocin and dopamine receptors show limited expression in the mouse lung. (**A**) Representative micrographs of mammary gland in lactating, and cabergoline treated lactating mice. Scale bars, 20 µm. (**B**) Normalized gene count of *Prlr* in lung epithelial (left) and endothelial (right) cells. **(C)** UMAP colored by log-normalized expression of broad cell-type marker genes: *Epcam* (epithelial), *Pecam1* (endothelial), *Ptprc* (immune), and *Col1a1* (mesenchymal), and prolactin receptor (*Prlr*), oxytocin receptor (*Oxtr*), and dopamine receptor (*Drd1a, Drd2, Drd3, Drd4*, and *Drd5*). Cells with zero detected counts are shown in grey. Color scale is shared across UMAPs and represents log-normalized expression clipped at the 99th percentile of expressing cells.

**Figure S6.**
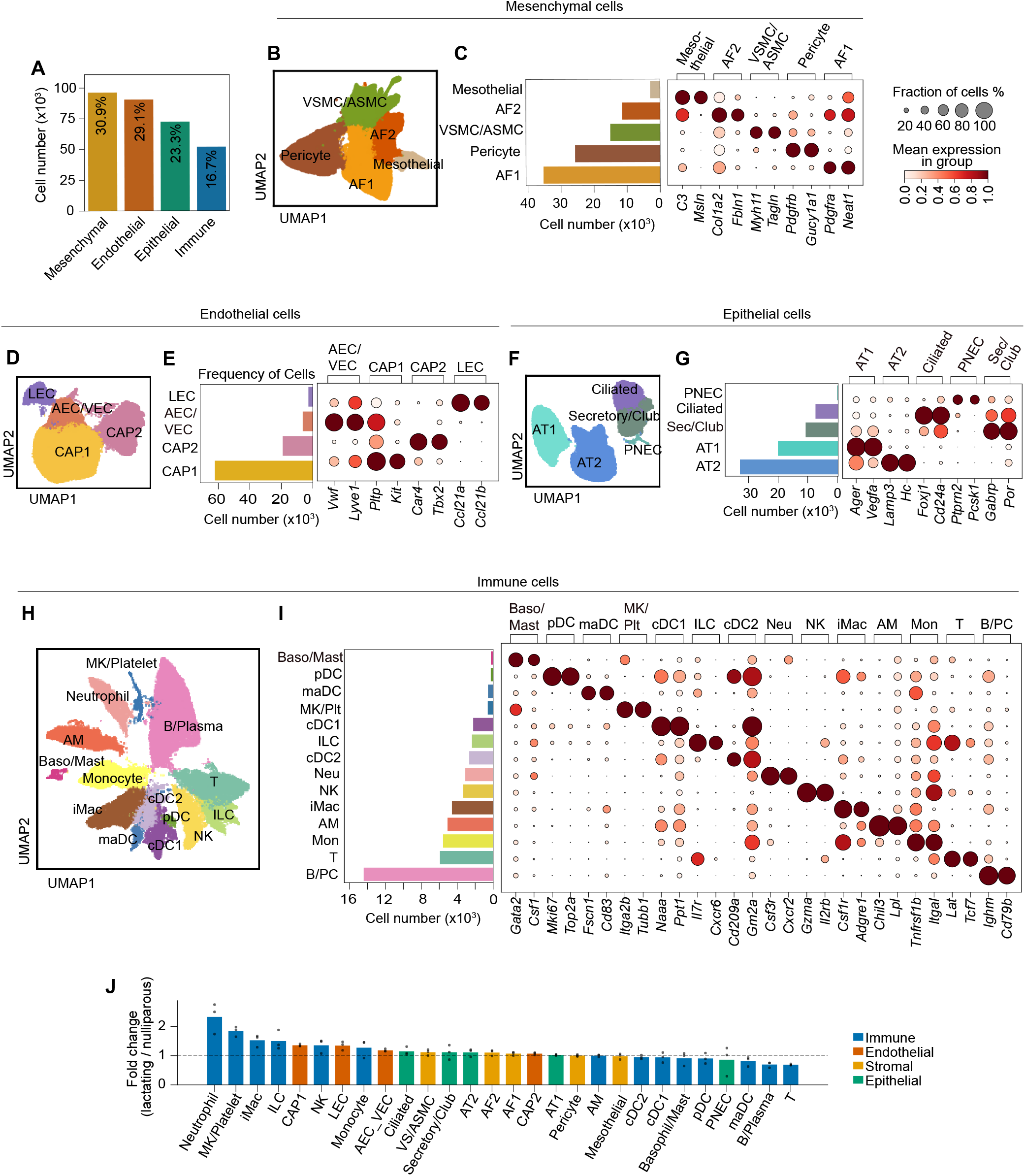
Spatial transcriptomic analysis of pulmonary remodeling. (**A**) Bar chart showing the percentage of cells per broad cell-type population. (**B**) UMAP representation of mesenchymal cells after independent reclustering, colored by subcluster identity. (**C**) Bar chart showing the number of cells per mesenchymal subcluster (left) and dot plot showing mean scaled expression of the top two marker genes per subcluster (right). Dot size indicates the fraction of cells expressing each gene; color indicates mean scaled expression. (**D**) UMAP representation of endothelial cells after independent reclustering, colored by subcluster identity. (**E**) As in (**C**), for endothelial subclusters. (**F**) UMAP representation of epithelial cells after independent reclustering, colored by subcluster identity. (**G**) As in (**C**), for epithelial subclusters. (**H**) UMAP representation of immune cells after independent reclustering, colored by subcluster identity. (**I**) As in (**C**), for immune subclusters. Neu, neutrophil; B/PC, B cell/plasma cell; Mon, monocyte; MK/Plt, megakaryocyte/platelet. (**J**) Bar chart showing fold change in cell density (cells per µm^2^; lactating versus nulliparous) for all clusters, colored by broad cell-type population. Dashed line indicates fold change = 1. Individual data points represent the fold change of each lactating replicate relative to mean nulliparous density.

**Figure S7.**
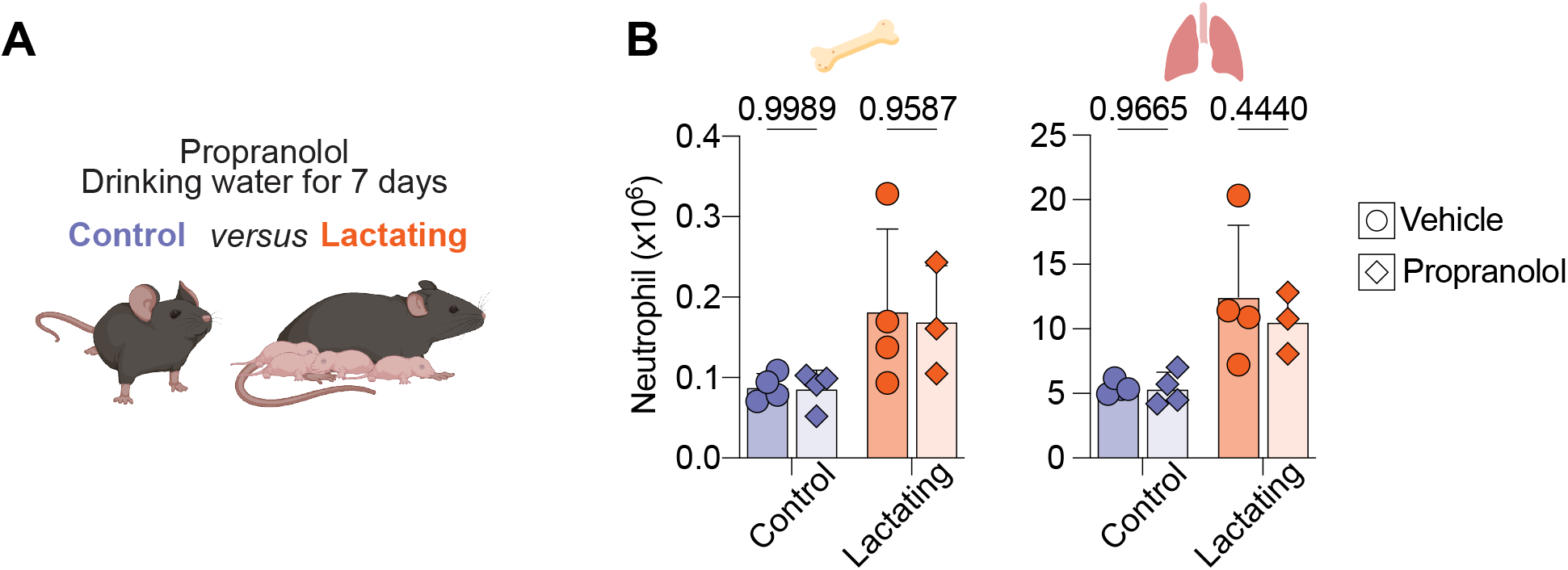
Limited contribution of β-adrenergic signaling to neutrophil accumulation. (**A**) Schematic of propranolol drinking water treatment. (**B**) Absolute cell number of bone marrow (left) and lung (right) neutrophils in vehicle and propranolol-treated control and lactating mice. Data in (B) represents mean ± SD and were analyzed by two-way ANOVA with Sidak’ s multiple comparisons correction. Each dot represents one animal, and *P* values are indicated.

## References

1. S. C. Ginther, H. Cameron, C. R. White, D. J. Marshall, Metabolic loads and the costs of metazoan reproduction. Science 384, 763–767 (2024).

2. D. Cansever, E. Petrova, S. Krishnarajah, C. Mussak, C. A. Welsh, W. Mildenberger, K. Mulder, V. Kreiner, E. Roussel, S. A. Stifter, M. Andreadou, P. Zwicky, N. P. Jurado, H. Rehrauer, G. Tan, Z. Liu, C. Blériot, F. Ronchi, A. J. Macpherson, F. Ginhoux, G. Natalucci, B. Becher, M. Greter, Lactation-associated macrophages exist in murine mammary tissue and human milk. Nat Immunol 24, 1098–1109 (2023).

3. C. A. Dawson, B. Pal, F. Vaillant, L. C. Gandolfo, Z. Liu, C. Bleriot, F. Ginhoux, G. K. Smyth, G. J. Lindeman, S. N. Mueller, A. C. Rios, J. E. Visvader, Tissue-resident ductal macrophages survey the mammary epithelium and facilitate tissue remodelling. Nat Cell Biol 22, 546–558 (2020).

4. S. K. Nyquist, P. Gao, T. K. J. Haining, M. R. Retchin, Y. Golan, R. S. Drake, K. Kolb, B. E. Mead, N. Ahituv, M. E. Martinez, A. K. Shalek, B. Berger, B. A. Goods, Cellular and transcriptional diversity over the course of human lactation. Proceedings of the National Academy of Sciences 119, e2121720119 (2022).

5. C. Hassel, B. Gausserès, L. Guzylack-Piriou, G. Foucras, Ductal Macrophages Predominate in the Immune Landscape of the Lactating Mammary Gland. Front. Immunol. 12 (2021).

6. A. Jaquish, E. Phung, X. Gong, P. Baldominos, S. Galván-Peña, I. Magill, I. Bursulaya, E. Marina, K. Bertrand, C. Chambers, A. R. Muñoz-Rojas, J. Agudo, D. Mathis, C. Benoist, D. Ramanan, Mammary intraepithelial lymphocytes and intestinal inputs shape T cell dynamics in lactogenesis. Nat Immunol 26, 1411–1422 (2025).

7. D. Corral, E. Ansaldo, J. Delaleu, A. C. Pichler, J. Kabat, C. Oguz, A. Teijeiro, D. Yong, M. Abid, C. A. Rivera, V. M. Link, K. Yang, L. Chi, J. Nie, O. Kamenyeva, Y. Fan, J. K. Y. Chan, F. Ginhoux, R. Bosselut, Y. Belkaid, Mammary intraepithelial lymphocytes promote lactogenesis and offspring fitness. Cell 188, 1662–1680.e24 (2025).

8. T. G. Wegmann, H. Lin, L. Guilbert, T. R. Mosmann, Bidirectional cytokine interactions in the maternal-fetal relationship: is successful pregnancy a TH2 phenomenon? Immunology Today 14, 353–356 (1993).

9. R. M. Samstein, S. Z. Josefowicz, A. Arvey, P. M. Treuting, A. Y. Rudensky, Extrathymic generation of regulatory T cells in placental mammals mitigates maternal-fetal conflict. Cell 150, 29–38 (2012).

10. D. A. Somerset, Y. Zheng, M. D. Kilby, D. M. Sansom, M. T. Drayson, Normal human pregnancy is associated with an elevation in the immune suppressive CD25+ CD4+ regulatory T-cell subset. Immunology 112, 38–43 (2004).

11. V. R. Aluvihare, M. Kallikourdis, A. G. Betz, Regulatory T cells mediate maternal tolerance to the fetus. Nat Immunol 5, 266–271 (2004).

12. T.-Y. Shao, J. M. Kinder, G. Harper, G. Pham, Y. Peng, J. Liu, E. J. Gregory, B. E. Sherman, Y. Wu, A. E. Iten, Y.-C. Hu, A. E. Russi, J. J. Erickson, H. Miller-Handley, S. S. Way, Reproductive outcomes after pregnancy-induced displacement of preexisting microchimeric cells. Science 381, 1324–1330 (2023).

13. C. Huang, A. Sun, J. Reyes, J. Ribeiro de Souza, T. R. Cafiero, K. H. Antunes Fernandes, F. M. S. Oliveira, Y. Qiao, P. Gazzinelli-Guimaraes, Y. Pritykin, A.I. Lim. Eosinophils drive intestinal remodelling and innate defence in reproduction. Nature 654, 209–218 (2026).

14. T. R. Martin, C. W. Frevert, Innate Immunity in the Lungs. Proc Am Thorac Soc 2, 403–411 (2005).

15. J. A. Whitsett, T. Alenghat, Respiratory epithelial cells orchestrate pulmonary innate immunity. Nat Immunol 16, 27–35 (2015).

16. R. J. Hewitt, C. M. Lloyd, Regulation of immune responses by the airway epithelial cell landscape. Nat Rev Immunol 21, 347–362 (2021).

17. A. Iwasaki, E. F. Foxman, R. D. Molony, Early local immune defenses in the respiratory tract. Nat Rev Immunol 17, 7–20 (2017).

18. J. Major, S. Crotta, K. Finsterbusch, P. Chakravarty, K. Shah, B. Frederico, R. D’Antuono M. Green, L. Meader, A. Suarez-Bonnet, S. Priestnall, B. Stockinger, A. Wack. Endothelial AHR activity prevents lung barrier disruption in viral infection. Nature, 621, 813–820 (2023).

19. C. H. Knight, E. Maltz, A. H. Docherty, Milk yield and composition in mice: effects of litter size and lactation number. Comp Biochem Physiol A Comp Physiol 84, 127–133 (1986).

20. H. Wu, A. Kuzmenko, S. Wan, L. Schaffer, A. Weiss, J. H. Fisher, K. S. Kim, F. X. McCormack, Surfactant proteins A and D inhibit the growth of Gram-negative bacteria by increasing membrane permeability. J Clin Invest 111, 1589–1602 (2003).

21. F. X. McCormack, J. A. Whitsett, The pulmonary collectins, SP-A and SP-D, orchestrate innate immunity in the lung. J Clin Invest 109, 707–712 (2002).

22. J. R. Wright, Immunoregulatory functions of surfactant proteins. Nat Rev Immunol 5, 58–68 (2005).

23. X. M. Bustamante-Marin, L. E. Ostrowski, Cilia and Mucociliary Clearance. Cold Spring Harb Perspect Biol 9, a028241 (2017).

24. K. C. Kim, Role of epithelial mucins during airway infection. Pulm Pharmacol Ther 25, 415–419 (2012).

25. P. Dhar, J. McAuley, The Role of the Cell Surface Mucin MUC1 as a Barrier to Infection and Regulator of Inflammation. Front. Cell. Infect. Microbiol. 9 (2019).

26. S. Devi, Y. Wang, W. K. Chew, R. Lima, N. A-González, C. N. Z. Mattar, S. Z. Chong, A. Schlitzer, N. Bakocevic, S. Chew, J. L. Keeble, C. C. Goh, J. L. Y. Li, M. Evrard, B. Malleret, A. Larbi, L. Renia, M. Haniffa, S. M. Tan, J. K. Y. Chan, K. Balabanian, T. Nagasawa, F. Bachelerie, A. Hidalgo, F. Ginhoux, P. Kubes, L. G. Ng, Neutrophil mobilization via plerixafor-mediated CXCR4 inhibition arises from lung demargination and blockade of neutrophil homing to the bone marrow. J Exp Med 210, 2321–2336 (2013).

27. A. B. Cohen, M. Rossi, Neutrophils in Normal Lungs. Am Rev Respir Dis 127, S3–S9 (1983).

28. C. Johansson, F. C. M. Kirsebom, Neutrophils in respiratory viral infections. Mucosal Immunol 14, 815–827 (2021).

29. K. G. Anderson, K. Mayer-Barber, H. Sung, L. Beura, B. R. James, J. J. Taylor, L. Qunaj, T. S. Griffith, V. Vezys, D. L. Barber, D. Masopust, Intravascular staining for discrimination of vascular and tissue leukocytes. Nat Protoc 9, 209–222 (2014).

30. I. Ballesteros, A. Rubio-Ponce, M. Genua, E. Lusito, I. Kwok, G. Fernández-Calvo, T. E. Khoyratty, E. van Grinsven, S. González-Hernández, J. Á. Nicolás-Ávila, T. Vicanolo, A. Maccataio, A. Benguría, J. L. Li, J. M. Adrover, A. Aroca-Crevillen, J. A. Quintana, S. Martín-Salamanca, F. Mayo, S. Ascher, G. Barbiera, O. Soehnlein, M. Gunzer, F. Ginhoux, F. Sánchez-Cabo, E. Nistal-Villán, C. Schulz, A. Dopazo, C. Reinhardt, I.A. Udalova, L. G. Ng, R. Ostuni, A. Hidalgo, Co-option of Neutrophil Fates by Tissue Environments. Cell 183, 1282–1297.e18 (2020).

31. D. Cerezo-Wallis, A. Rubio-Ponce, M. Richter, E. Pitino, I. Kwok, G. Marteletto, A. C. Guanolema-Coba, C. Shih, R.-K. Huang, A. Moraga, N. B. Bravo, S. Doré, S. Callejas, D. G. Aragonés, D. Jiménez-Carretero, D. Martin, S. Ovadia, T. Vicanolo, G. Crainiciuc, J. Sicilia, T. Deng, A. Martin, J. Zhang, M. I. Cuartero, D. M. Giraldo, A. Garcia-Culebras, A. Aroca-Crevillen, S. Martín-Salamanca, C. Torroja, M. Ruiz, I. Ruano, M.S. F. Ng, J. Hou, Y. Wang, M. Zhang, J. Pu, A. Herruzo, D. C. van Oordt, S. Chang, A. E. Downie, F. Chen, A. L. Graham, W. C. Gause, P. O. Fiset, J. D. Spicer, H. Heyn, M. A. Zuriaga, J. A. Bernal, I. A. Udalova, M. A. Moro, K. de Bock, A. Dopazo, J. J. Fuster, F. Sánchez-Cabo, J. C. Nieto, G. F. Calvo, J. Skokowa, O. Soehnlein, D. F. Quail, L. A. Walsh, L. G. Ng, A. Hidalgo, I. Ballesteros, Architecture of the neutrophil compartment. Nature 649, 1003–1012 (2026).

32. L. G. Ng, I. Ballesteros, M. A. Cassatella, M. Egeblad, Z. G. Fridlender, D. Gabrilovich, Q. Gao, Z. Granot, R. Grieshaber-Bouyer, H. L. Grimes, C. C. Hedrick, A. Hidalgo, M. J. Kaplan, P. Kubes, G. S. Ling, L. Lu, H. R. Luo, T. N. Mayadas, N. M. Moutsopoulos, M. Ng, P. A. Nigrovic, R. Ostuni, M. J. Pittet, D. F. Quail, C. Silvestre-Roig, O. Soehnlein, I. A. Udalova, R. Xue, N. Zhang, I. Kwok, From complexity to consensus: A roadmap for neutrophil classification. Immunity 58, 1890–1903 (2025).

33. M. Nagai, M. Moriyama, C. Ishii, H. Mori, H. Watanabe, T. Nakahara, T. Yamada, D. Ishikawa, T. Ishikawa, A. Hirayama, I. Kimura, A. Nagahara, T. Naito, S. Fukuda, T. Ichinohe, High body temperature increases gut microbiota-dependent host resistance to influenza A virus and SARS-CoV-2 infection. Nat Commun 14, 3863 (2023).

34. H. Lei, Y. Xu, H. Zhang, B. Zhang, W. Luo, X. Liu, H. Zhang, J. Yang, W. Wen, P. Wang, S. Xu, Multiomics analysis unveils key biomarkers during dynamic progress of IAV infection in mice. Front. Immunol. 16 (2025).

35. T. Madan, P. Eggleton, U. Kishore, P. Strong, S. S. Aggrawal, P. U. Sarma, K. B. Reid, Binding of pulmonary surfactant proteins A and D to Aspergillus fumigatus conidia enhances phagocytosis and killing by human neutrophils and alveolar macrophages. Infect Immun 65, 3171–3179 (1997).

36. M. M. Mircescu, L. Lipuma, N. van Rooijen, E. G. Pamer, T. M. Hohl, Essential role for neutrophils but not alveolar macrophages at early time points following Aspergillus fumigatus infection. J Infect Dis 200, 647–656 (2009).

37. M. Onji, V. Sigl, T. Lendl, M. Novatchkova, A. Ullate-Agote, A. Andersson-Rolf, I. Kozieradzki, R. Koglgruber, T.-P. Pai, D. Lichtscheidl, K. Nayak, M. Zilbauer, N. A. Carranza García, L. K. Sievers, M. Falk-Paulsen, S. J. F. Cronin, A. Hagelkruys, S. Sawa, L. C. Osborne, P. Rosenstiel, M. Pasparakis, J. Ruland, H. Takayanagi, H. Clevers, B.-K. Koo, J. M. Penninger, RANK drives structured intestinal epithelial expansion during pregnancy. Nature 637, 156–166 (2025).

38. M. Guo, M. P. Morley, C. Jiang, Y. Wu, G. Li, Y. Du, S. Zhao, A. Wagner, A. C. Cakar, M. Kouril, K. Jin, N. Gaddis, J. A. Kitzmiller, K. Stewart, M. C. Basil, S. M. Lin, Y. Ying, A. Babu, K. A. Wikenheiser-Brokamp, K. S. Mun, A. P. Naren, G. Clair, J. N. Adkins, G. S. Pryhuber, R. S. Misra, B. J. Aronow, T. L. Tickle, N. Salomonis, X. Sun, E. E. Morrisey, J. A. Whitsett, S. Lin, Y. Xu, Guided construction of single cell reference for human and mouse lung. Nat Commun 14, 4566 (2023).

39. H. Zhao, X. Ma, P. Chen, B. Liu, J. Wei, J. Zhang, A. Desai, A. L. Frump, O. Rafikova, M. B. Fallon, S. Qiu, Z. Dai, Enhanced Spatial Transcriptomics Analysis of Mouse Lung Tissues Reveals Cell-Specific Gene Expression Changes Associated with Pulmonary Hypertension. J Respir Biol Transl Med 2, 10004 (2025).

40. T. Liu, L. M. Miller, B. P. Dresden, C. Zhao, Y. Pillai, A. Rangel-Garcia, M. de la L. Garcia-Hernandez, J. Rangel-Moreno, W. Chen, J. F. Alcorn, Influenza virus and Staphylococcus aureus super-infection disrupts spatially coordinated cellular immunity in the mouse lung. Commun Biol 8, 1837 (2025).

41. A. Scanzano, M. Cosentino, Adrenergic regulation of innate immunity: a review. Front Pharmacol 6, 171 (2015).

42. T. Liu, L. Yang, X. Han, X. Ding, J. Li, J. Yang, Local sympathetic innervations modulate the lung innate immune responses. Science Advances 6, eaay1497 (2020).

43. B. M. Assas, J. I. Pennock, J. A. Miyan, Calcitonin gene-related peptide is a key neurotransmitter in the neuro-immune axis. Front Neurosci 8, 23 (2014).

44. P. Baral, B. D. Umans, L. Li, A. Wallrapp, M. Bist, T. Kirschbaum, Y. Wei, Y. Zhou, V. K. Kuchroo, P. R. Burkett, B. G. Yipp, S. D. Liberles, I. M. Chiu, Nociceptor sensory neurons suppress neutrophil and γδ T cell responses in bacterial lung infections and lethal pneumonia. Nat Med 24, 417–426 (2018).

45. K. J. Blake, X. R. Jiang, I. M. Chiu, Neuronal Regulation of Immunity in the Skin and Lungs. Trends in Neurosciences 42, 537–551 (2019).

46. A. M. Ehlers, I. M. Guerrero-Fonseca, C. Altier, B. G. Yipp, S. Talbot, Wired for immunity: neuroimmune control of the lung by sensory neurons. Nat. Rev. Neurosci. 27, 481–493 (2026).

47. T. E. Williamson, S. L. Brusatte, G. P. Wilson, The origin and early evolution of metatherian mammals: the Cretaceous record. Zookeys, 1–76 (2014).

48. M. dos Reis, J. Inoue, M. Hasegawa, R. J. Asher, P. C. J. Donoghue, Z. Yang, Phylogenomic datasets provide both precision and accuracy in estimating the timescale of placental mammal phylogeny. Proc Biol Sci 279, 3491–3500 (2012).

49. J. E. Tarver, M. Dos Reis, S. Mirarab, R. J. Moran, S. Parker, J. E. O’ Reilly B. L. King, M. J. O’Connell R. J. Asher, T. Warnow, K. J. Peterson, P. C. J. Donoghue, D. Pisani, The Interrelationships of Placental Mammals and the Limits of Phylogenetic Inference. Genome Biol Evol 8, 330–344 (2016).

50. S. Denk, R. P. Taylor, R. Wiegner, E. M. Cook, M. A. Lindorfer, K. Pfeiffer, S. Paschke, T. Eiseler, M. Weiss, E. Barth, J. D. Lambris, M. Kalbitz, T. Martin, H. Barth, D. A. C. Messerer, F. Gebhard, M. S. Huber-Lang, Complement C5a-induced changes in neutrophil morphology during inflammation. Scand J Immunol 86, 143–155 (2017).

51. E. van Grinsven, A. K. Mukherjee, I. A. Udalova, Shaping neutrophil morphology and function: the importance of a segmented nucleus. Nat Rev Immunol 26, 269–283 (2026).

52. K. Branchfield, L. Nantie, J. M. Verheyden, P. Sui, M. D. Wienhold, X. Sun, Pulmonary neuroendocrine cells function as airway sensors to control lung immune response. Science 351, 707–710 (2016).

53. P. Sui, D. L. Wiesner, J. Xu, Y. Zhang, J. Lee, S. Van Dyken, A. Lashua, C. Yu, B. S. Klein, R. M. Locksley,G. Deutsch, X. Sun, Pulmonary neuroendocrine cells amplify allergic asthma responses. Science 360, eaan8546 (2018).

54. H. Song, E. Yao, C. Lin, R. Gacayan, M.-H. Chen, P.-T. Chuang, Functional characterization of pulmonary neuroendocrine cells in lung development, injury, and tumorigenesis. Proceedings of the National Academy of Sciences 109, 17531–17536 (2012).

55. C. Y. Feigin, J. A. Moreno, R. Ramos, S. A. Mereby, A. Alivisatos, W. Wang, R. van Amerongen, J. Camacho, J. J. Rasweiler, R. R. Behringer, B. Ostrow, M. V. Plikus, R. Mallarino, Convergent deployment of ancestral functions during the evolution of mammalian flight membranes. Science Advances 9, eade7511 (2023).

56. J. Niu, M. Cui, X. Yang, J. Li, Y. Yao, Q. Guo, A. Lu, X. Qi, D. Zhou, C. Zhang, L. Zhao, G. Meng, Microbiota-derived acetate enhances host antiviral response via NLRP3. Nat Commun 14, 642 (2023).

57. J. V. Desai, M. A. Zarakas, A. L. Wishart, M. Roschewski, M. A. Aufiero, A. Donkò, G. Wigerblad, N. Shlezinger, M. Plate, M. R. James, J. K. Lim, G. Uzel, J. R. Bergerson, I. Fuss, R. A. Cramer, L. M. Franco, E. S. Clark, W. N. Khan, D. Yamanaka, G. Chamilos, J. El-Benna, M. J. Kaplan, L. M. Staudt, T. L. Leto, S. M. Holland, W. H. Wilson, T. M. Hohl, M. S. Lionakis, BTK drives neutrophil activation for sterilizing antifungal immunity. J Clin Invest 134, e176142 (2024).

58. K. Shimizu, N. P. Keller, Genetic involvement of a cAMP-dependent protein kinase in a G protein signaling pathway regulating morphological and chemical transitions in Aspergillus nidulans. Genetics 157, 591–600 (2001).

